# The repertoire of resistance mutations selected by a *Pseudomonas aeruginosa* type IV pilus-targeting lytic bacteriophage

**DOI:** 10.1101/2025.11.17.688846

**Authors:** Veronica N. Tran, Hanjeong Harvey, Tanisha S. Lahane, Lori L. Burrows

## Abstract

*P. aeruginosa* type IV pili (T4P) are complex surface-exposed nanomachines used for twitching motility and biofilm formation. Exposure to bacteriophages that use T4P as their primary receptor can select for resistance mutations that reduce or abolish pilus expression. Since T4P are important virulence factors, such loss can affect pathogenicity and fitness, desirable outcomes for phage therapy. Here, we evaluated the repertoire of mutations in *P. aeruginosa* PAO1 exposed to the pilus-specific lytic phage PO4 and their ability to revert after removal of phage. Resistant isolates ranged from hyperpiliated to bald, but none retained twitching motility. We focused on isolates that still expressed surface pili, including one with an internal 4-amino acid duplication in the essential prepilin peptidase/methyltransferase, PilD. This duplication, distal to the predicted active sites, delayed prepilin processing. Accumulation of unprocessed subunits suppressed expression of new prepilins via inhibition of PilSR two-component system activity, restricting availability of functional subunits and conferring phage resistance. Introduction of a PilS point mutation that makes cells insensitive to accumulation of unprocessed pilins restored motility and phage susceptibility. Of the mutants tested, only those with a duplication in *pilD* recovered wild-type motility following removal of phage pressure, likely through slip-strand mispairing as a codon-modified variant did not revert. This work shows that resistance to pilus-specific phages does not require loss of pilus expression, and certain mutations can allow bacteria to regain pilus function. Characterizing spontaeous mutations selected by phages can help to define the function and vulnerabilities of the type IV pilus system.

**SIGNIFICANCE:** As use of phages to treat antibiotic-resistant pathogens such as *Pseudomonas aeruginosa* increases, it is important to understand potential outcomes of phage exposure. Most therapeutic *P. aeruginosa* phages use lipopolysaccharides (LPS) or T4P as primary receptors. Studying the properties of strains resistant to T4P-targeting phages can guide the design of phage cocktails to mitigate treatment resistance. We show that depending on the mutation, some phage-resistant strains can revert to wild-type sequences, emphasizing the importance of combining diverse phages to supress resurgence. By characterizing mutations that confer resistance, we can better understand whether pilus structural or regulatory components are more likely to be lost. Using phages to select for loss of pilus function represents an unbiased approach to identify new mutations in pilus-related proteins, shedding light on understudied components. Building a database of such mutations will help guide strategies to target and disarm this key *P. aeruginosa* virulence factor.

## INTRODUCTION

Type IV pili (T4P) are the most common type of prokaryotic surface adhesin and are involved in motility, surface sensing, and DNA uptake (1–3). These hair-like filaments allow for non-specific attachment of bacterial pathogens to surfaces and contribute to establishment of infection and dissemination of pathogens in plant and animal hosts (4–6). They also serve as receptors for select bacteriophages (phages) that bind to the pili to initiate their infection cycle (7–9). Here we investigated the effects of exposing *Pseudomonas aeruginosa* to a T4P-targeting phage, focusing on mechanisms of resistance and potential for reversion to wild-type phenotypes following removal of the phage.

*P. aeruginosa* is an opportunistic antibiotic-resistant pathogen considered by the World Health Organization to be a top priority for development of new therapies, and a model species for the study of T4P biology (3, 10). There are 3 major subfamilies of T4P: T4aP, T4bP, and T4cP (Tad), and most *P. aeruginosa* strains express T4aP (2, 11). T4aP are repeatedly and rapidly extended and retracted by a complex nanomachine that spans the entire cell envelope (**Figure 1**). Major pilins (PilA) are lollipop-shaped subunits that make up the length of the pilus, while the minor pilin complex, consisting of minor pilins FimU, PilVWX and PilE, plus the non-pilus adhesin PilY1, is located at the tip (3, 12). Both major and minor pilins are initially expressed as inner membrane-embedded prepilins, with a short, positively charged leader sequence that is cleaved on the cytoplasmic side of the membrane by the prepilin peptidase PilD, followed by methylation of the new N-terminus, also by PilD (13, 14). Maturation of pilins by removal of the leader sequence is essential to make them competent for assembly, while methylation is not (15, 16).

**Figure 1.**
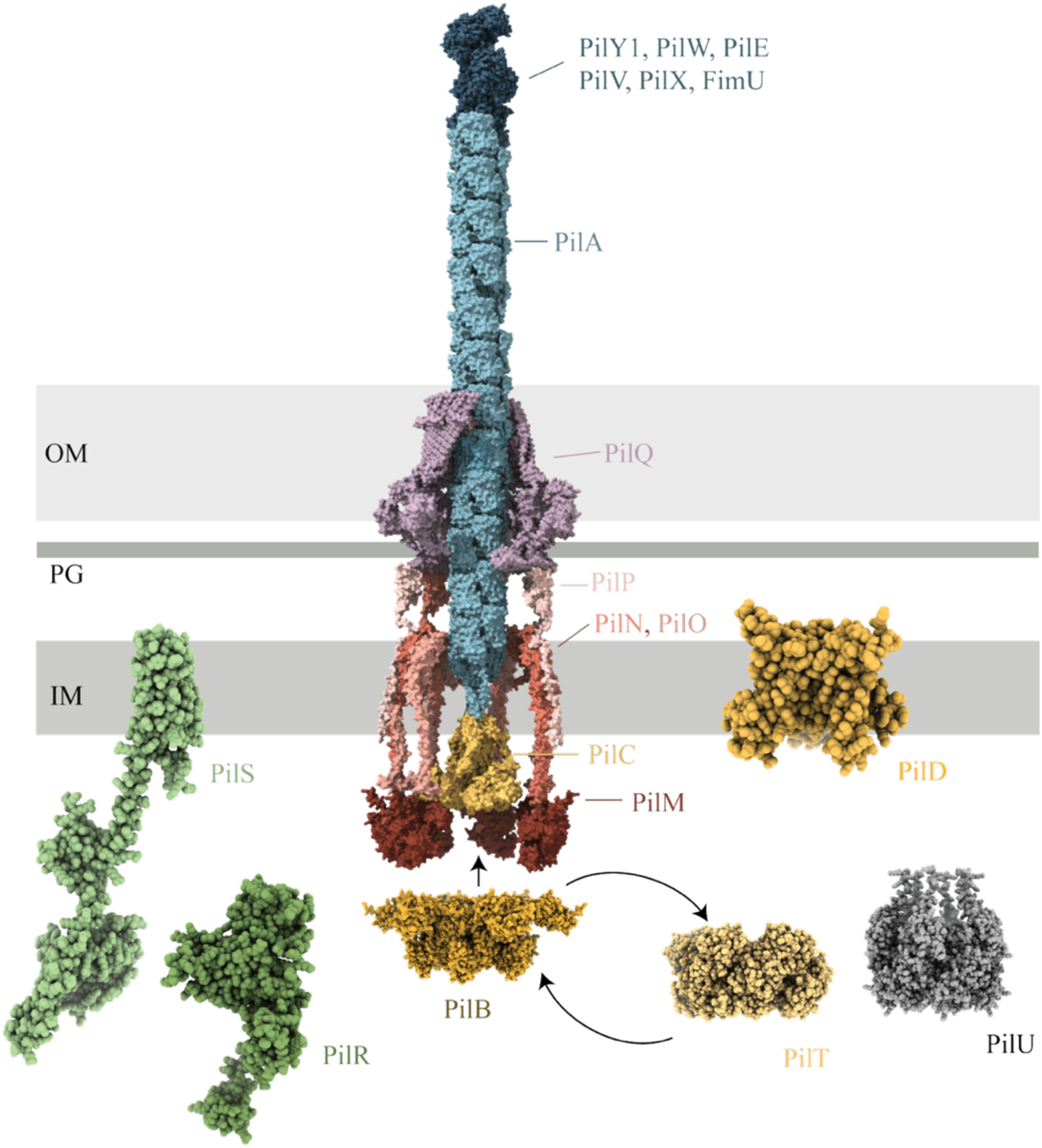
Structural models of the type IV pilus machinery. The type IV pilus is assembled by a complex protein nanomachine (EMD-43426) composed of four subcomplexes. The outer membrane secretin (PilQ) is shown in purple, the alignment subcomplex (PilM, PilN, PilO, PilP) in red, inner membrane motor subcomplex (PilC, PilB [AlphaFold3], PilT [PDB: 3JVU], PilU [grey, AlphaFold3], PilD [AlphaFold3]) in yellow and orange. PilB and PilT both interact with PilC, but not simultaneously, as indicated by the alternating arrows.The pilus subcomplex (major pilin subunits PilA and minor pilin subcomplex PilY1, PilE, PilW, PilV, PilX, FimU [AlphaFold3]) are shown in blue. The PilS-PilR two component system (green, AlphaFold3) regulates *pilA* expression. Coloured subcomplexes and labels indicate components in which we identified mutations conferring phage resistance. AlphaFold3 model confidence information is available in **Supplementary** Figure 1. Models are not to scale.

After formation of a minor pilin tip complex in the cytoplasmic membrane, mature pilins are added sequentially from inner membrane pools to the base of the complex, and the pilus is thus extended through the periplasm and out of the cell (3). Retraction reverses this process, disassembling the filament and returning the major pilins to the membrane to be used for subsequent rounds of extension (3, 17). Extension and retraction are powered by the extension ATPase PilB and retraction ATPases PilT and PilU, respectively, and coordinated by the platform protein PilC (18–22). The inner-membrane alignment subcomplex (PilMNOP) and the outer membrane secretin channel (PilQ) form a gated conduit that guides the growing pilus through the cell envelope (23–26).

Pilus extension and retraction facilitates a form of movement on surfaces called twitching motility (27), and twitching-deficient cells have reduced virulence, impaired ability to infect, and form abnormal biofilm structures (3, 28). Phages that use the pilus as a primary receptor take advantage of its retraction to gain access to the cell surface, where they may interact with secondary receptors and inject their genetic material (18, 29, 30). Such phages may bind to the major, or more rarely, the minor pilins (9, 31) although the specific molecular details of recognition and binding are only beginning to emerge (32).

Phages and bacteria are in an ongoing arms race, with each adapting as the other evolves new means of escape. Bacterial resistance mechanisms act at all stages of phage infection, including initial phage adsorption, which is thought to occur stochastically (33). Phage receptors can be mutated, post-translationally modified, masked by polysaccharides, or their surface chemistry or length altered to avoid phage infection (9, 34, 35). These changes can have implications for bacterial fitness, virulence, and recognition by the immune system, as well as the design of interventions. With antibiotic resistance on the rise, phage therapy is being revisited for treatment of persistent infections (36). Therefore, it is important to understand how bacteria might respond to and evade phage infection through surface modifications. In a clinical context, T4P-targeting phages could be used to steer infecting strains towards reduced fitness and/or increased susceptibility to treatment through loss of T4P function (37, 38). However, selection for mutations that could easily revert (e.g. phase variants or sequence duplications) or that could be bypassed by suppressors may allow for return to virulence once phage therapy is completed.

Here we examined the repertoire and potential reversibility of mutations conferring resistance following exposure of *P. aeruginosa* PAO1 to PO4, a T4P-targeting lytic phage (7). Phage-resistant mutants (PRMs) were sequenced to identify relevant mutations, and the ability of those mutations to confer cross-resistance to other phages tested. Unique resistance mutations in multiple regulatory and structural genes were identified, including a novel duplication in the essential prepilin peptidase PilD that led to protracted delays in cleavage of its prepilin substrates. Since most of the mutations identified were point mutations, small insertions/deletions (indels) or short sequence duplications, adaptation in the absence of phage pressure was evaluated. This analysis provided a snapshot of the range and stability of phage-selected T4aP mutations, important information for future design of phage cocktails.

## RESULTS

### T4aP-targeting phage PO4 selects for mutations in a variety of T4aP genes

Phage PO4 is a lytic phiKMV-like T4P-targeting phage that infects *P. aeruginosa* strain PAO1 (7). We used PAO1 for these studies because it lacks a functional CRISPR system, therefore it relies on resistance mechanisms that act at other steps of phage infection, such as receptor recognition. Following co-culture of PAO1 with PO4, 128 PO4-resistant colonies of *P. aeruginosa* were recovered. None of the phage-resistant mutants (PRMs) twitched in a standard agar sub-surface assay (**Supplementary Figure S2**) suggesting that these strains had non-functional T4aP (3, 27).

Whole genome sequencing followed by breseq analysis was used to identify relevant mutations. Among the 128 PRMs, we identified 28 unique mutations spanning 16 different pilus-related genes (Figure 1 and **Table 1**), suggesting that some mutants were clonal. Seventeen mutations resulted in truncation of the gene product and predicted loss of the protein. The remainder were small indels or single nucleotide polymorphisms (SNPs) predicted to alter protein function without disrupting expression. Most mutations affected only single genes, but one PRM had a large 7559-base deletion that spanned *pilS-pilY1*, removing a region that includes most of the minor pilin operon, plus the *pilSR* genes encoding a two-component system (TCS) that regulates PilA expression (39). Mutations spanned all the T4P structural subcomplexes (pilus, motor, alignment and secretin) plus PilS-PilR. Multiple non-clonal mutations in some genes were identified, including *pilM* (4 different PRMs), 3 in *pilB*, 3 in *pilR*, and 2 in each of *pilA*, *pilC*, *pilD*, and *pilQ*. There was no correlation between the size of the gene and the number of unique mutations identified. For example, only one mutation in *pilY1* (3486 bp) was recovered even though it is nearly double the length of *pilB,* in which multiple mutations were identified.

**Table 1.**
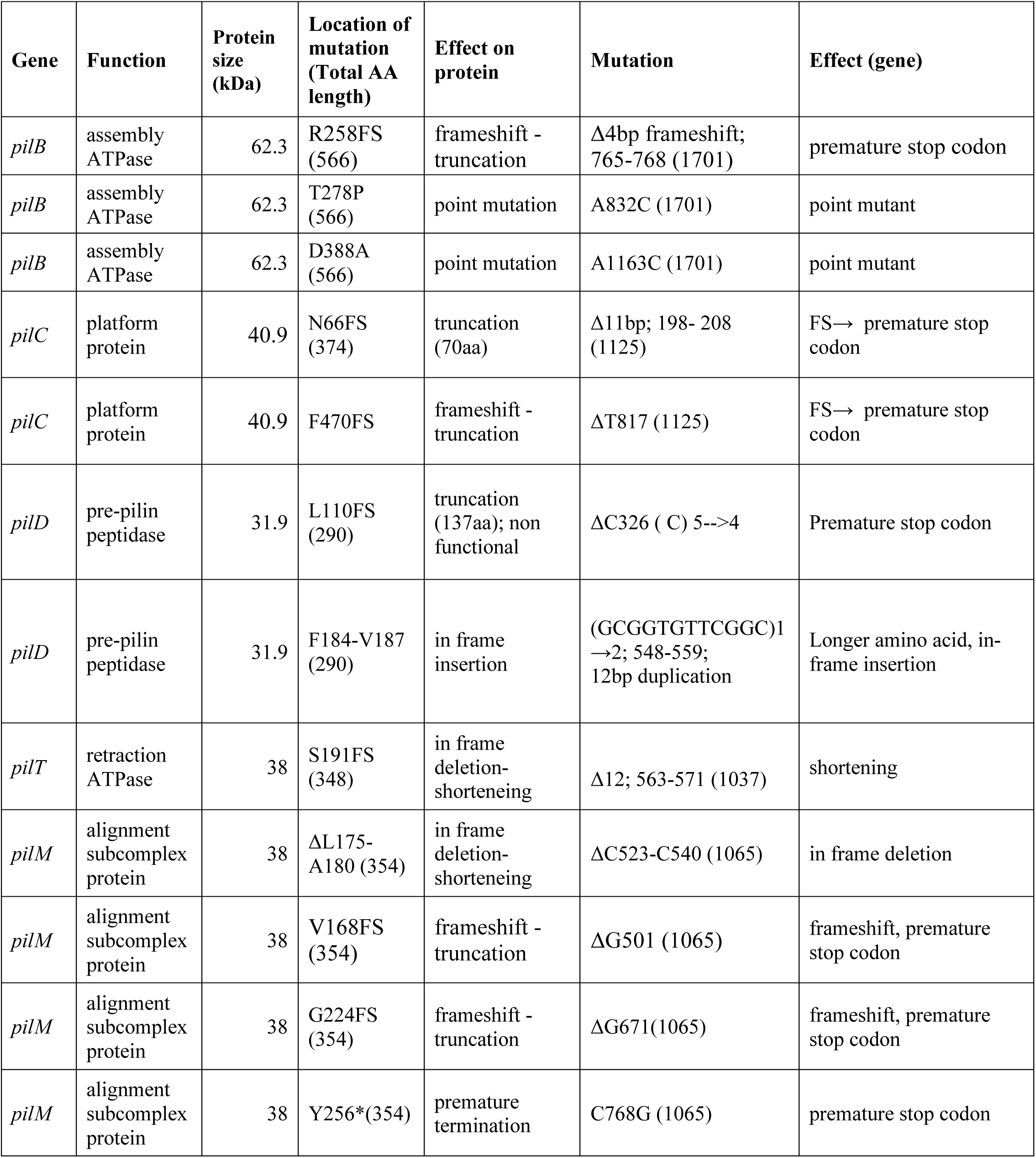
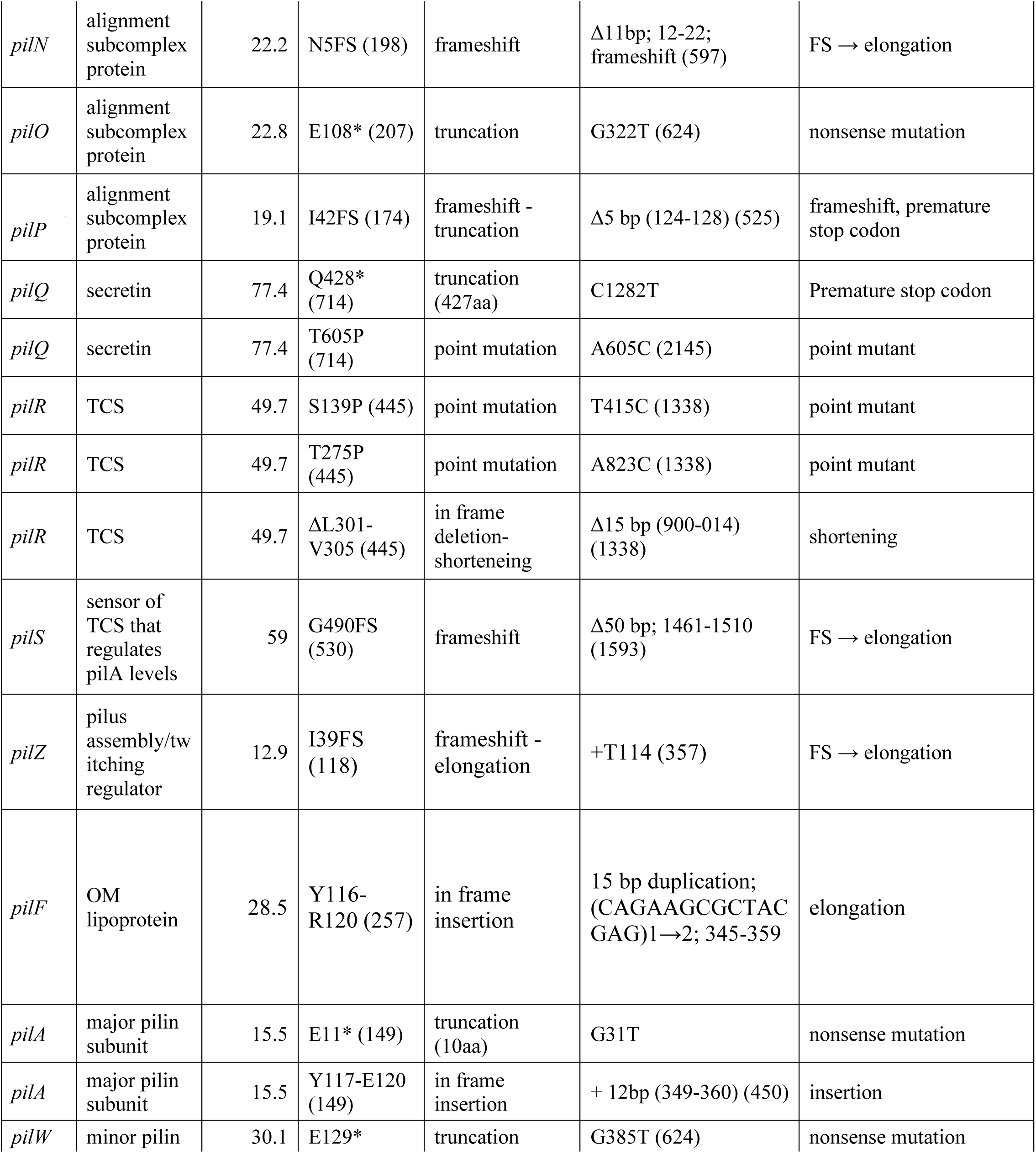
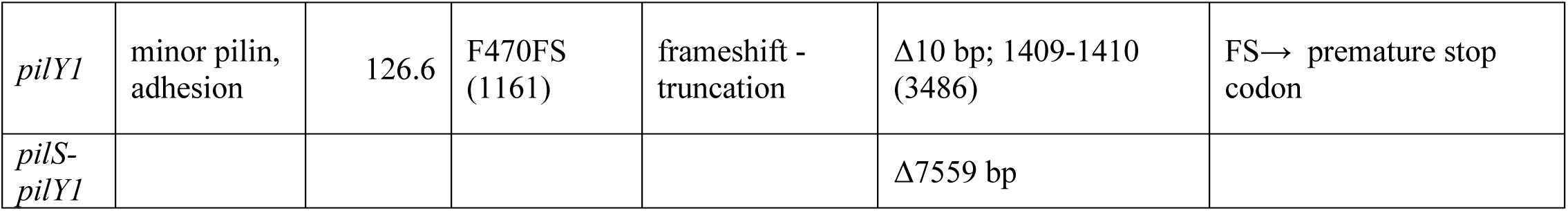
Mutations identified in Phage Resistant Mutants (PRMs)

Intriguingly, in four of the PRMs we could identify no mutations, despite repeated sequencing, pooling of the sequence data, and careful analysis of read depth (below). These strains remained phage insensitive after passage, suggesting they are stably resistant descendants of the PAO1 parent strain.

While complete loss of a gene product may result in an expected phenotype, SNPs and indels in the PRMs could generate informative intermediate phenotypes. Therefore, the levels of intracellular and extracellular pilins were assessed to test whether the mutations we identified resulted in loss of pilin expression and/or pilus assembly. Most PO4 PRMs expressed the major pilin subunit PilA at wild-type levels in whole cells, but failed to produce extracellular pili **(Supplementary Figure S2**). These data indicated that phage resistance in these PRMs results from impaired function of the pilus assembly machinery (**Table 1, Supplementary Figure S2**).

We identified three PRMs with recoverable extracellular pili, despite their lack of motility (Figure 2). One of these had a 4-base deletion (residues 568-571) in *pilT* encoding the retraction ATPase that caused a frameshift leading to a premature stop codon. *pilT* mutants produce extracellular pili, but PilT-mediated pilus retraction is required for PO4 susceptibility. Thus, loss of this protein is consistent with previously reported phage susceptibility patterns (18). The other two mutations were in the extension ATPase PilB and the prepilin peptidase/methyltransferase PilD. Since mutations in those proteins have not been reported to confer phage resistance while maintaining pilus expression, we examined them more closely.

**Figure 2.**
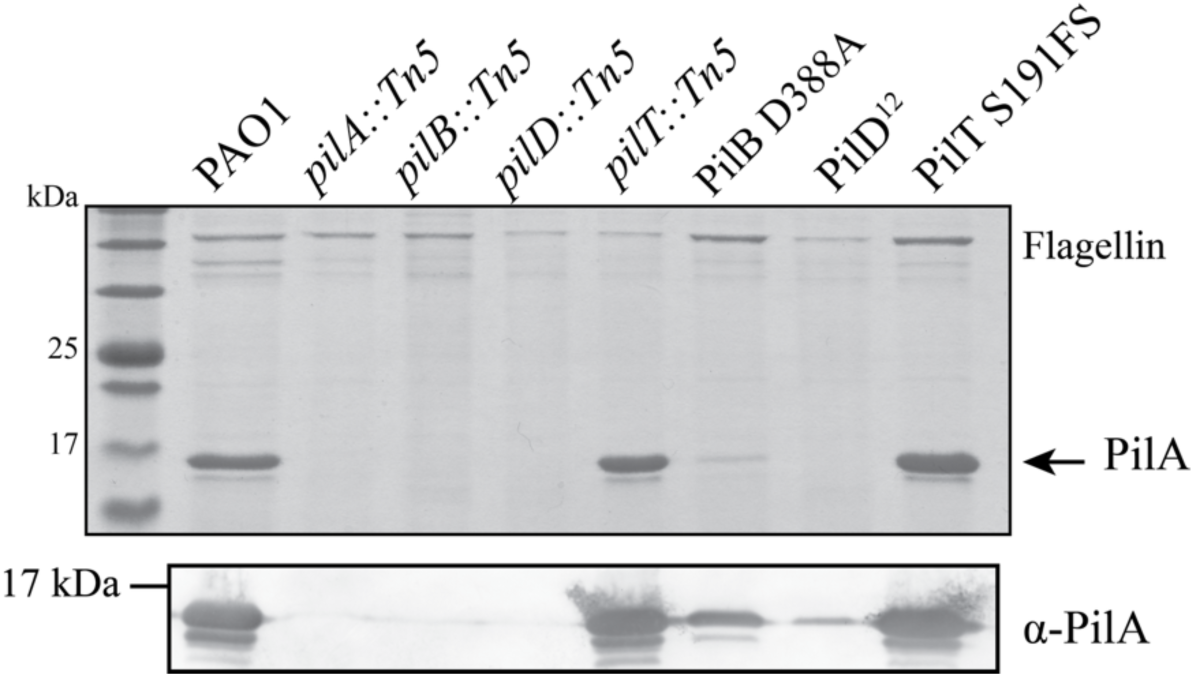
Phage-resistant mutants with lesions in PilB, PilD, or PilT still had recoverable surface pili. Sheared surface protein samples separated on 15% Coomassie-stained SDS-PA gel (top) and Western immunoblot using α-PilA antisera (bottom) of sheared surface protein samples show pilins can be recovered in these phage resistant mutants. Blot is representative of three independent experiments.

### PilB D388A confers phage resistance without loss of pilus extension

The assembly ATPase PilB is a member of the Additional Strand Catalytic ‘E’ superfamily (40). Its Walker B motif (residues D390-E396) coordinates a Mg^2+^ required for ATP hydrolysis (40–42), and this activity is essential for pilus assembly. We identified a PilB D388A mutation adjacent to the Walker B motif. Western blotting of whole cell lysates with a polyclonal PilB antibody showed that PilB D388A was expressed at levels similar to wild type (Figure 3A) suggesting the mutation does not affect protein stability, even though the mutant had fewer recoverable surface pili than wild type (Figure 2**, Supplementary Figure S1**). To test whether PilB D388A confers resistance to other T4P-targeting phages, the mutant was challenged with broad host range temperate phage DMS3 (43) and JBD68, which uses a minor pilin as its receptor (44) (Figure 3B). The PRM was resistant to both those T4P phages.

**Figure 3.**
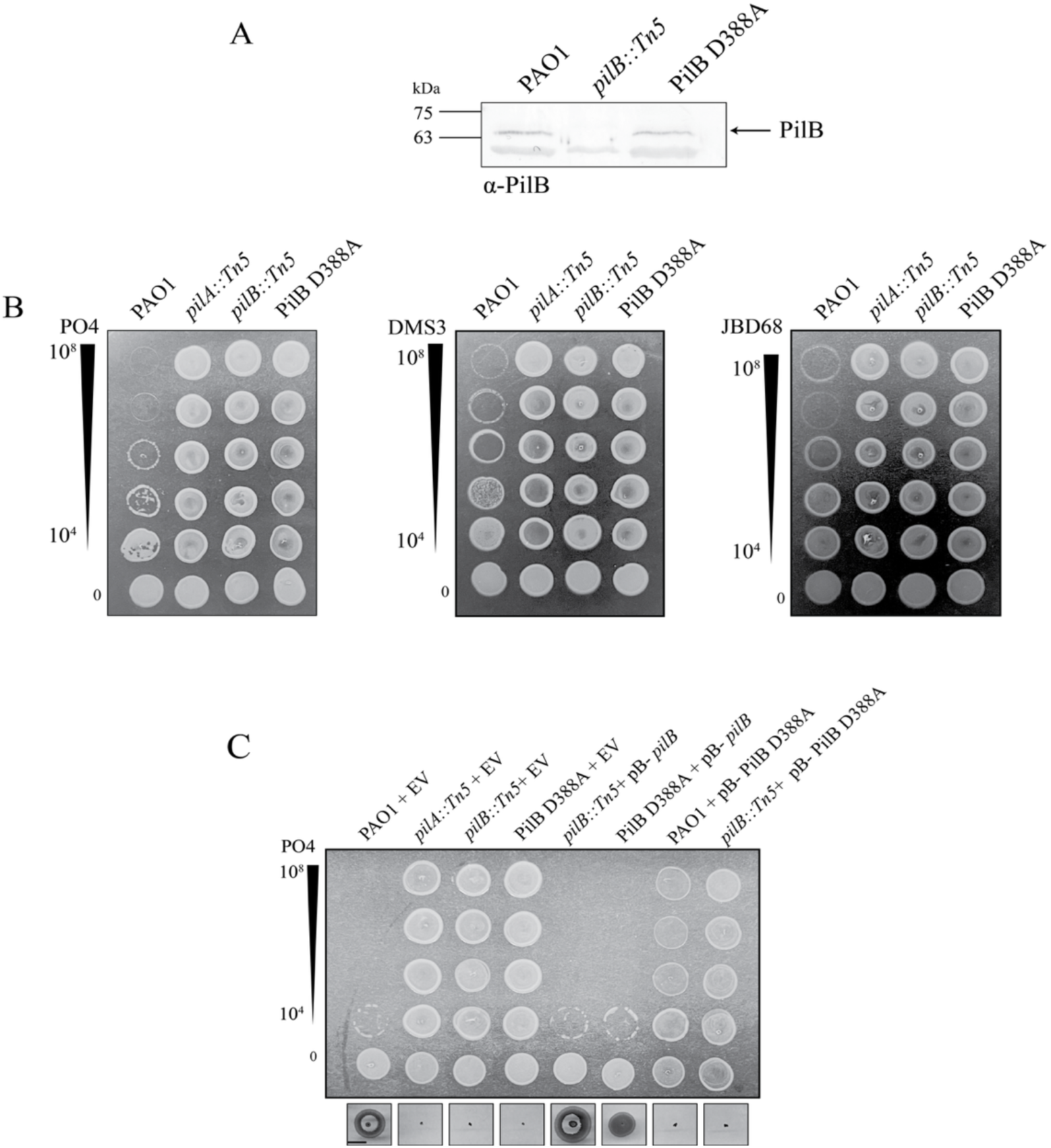
PilB D388A is stable and confers resistance to multiple T4P-targeting phages. **A)** Western immunoblot of whole cell lysates using α-PilB antisera showed that PilB D388A levels are comparable to wild-type PilB. Blot is representative of three independent experiments. **B)** PilB D338A is resistant to three different T4P-targeting phages. Serially diluted phages PO4, DMS3, and JBD68 were combined with standardized bacteria liquid cultures were spotted on 0.6% LB agar. Phage titres (PFU) are indicated to the left. **C)** Overexpression of *pilB in trans* in PilB D388A rescues phage susceptibility and twitching motility. Complementation of PAO1 or a *pilB::Tn5* mutant with PilB D388A abolishes twitching motility and phage susceptibility. Samples were induced using 0.2% L-arabinose. Scale bar indicates 1 cm. EV= empy vector, pB= pBADGr. Images are representative of three independent experiments.

Since PilB functions as a homohexamer (40), we tested for possible dominant-negative effects of expressing the mutant allele *in trans*. When wild-type PilB was expressed from a plasmid in PilB D388A, phage susceptibility and twitching motility were recovered, although motility was approximately 75% of wild-type (Figure 3C). In the reciprocal experiment, expression of PilB D388A from a multicopy plasmid in the wild type abolished twitching motility and phage susceptibility (Figure 3C). Similarly, complementing PilB-deficient strains with PilB D388A failed to restore twitching motility or phage susceptibility (Figure 3C), providing further evidence that this mutation impairs pilus function.

### A partial PilD duplication delays pilin maturation to confer phage resistance

The third PRM with recoverable surface pili had a 12-base duplication in *pilD* (PilD^12^; **Table 1**, Figure 2). PilD is a membrane-embedded bifunctional prepilin peptidase that cleaves the type III signal sequence of both T4P pilins and type II secretion system (T2SS) endopilins, and methylates the first residue of the mature (endo)pilins (14, 45–47). The duplication added four residues (duplication of F184-V187) to a helix that is distal to the predicted peptidase and methyltransferase active sites (Figure 4A). PRMs with this mutation had few recoverable extracellular pili that were not visible on SDS-PAGE, although they could be detected on Western blots (Figure 2), and did not twitch (**Supplementary Figure S2**).

**Figure 4.**
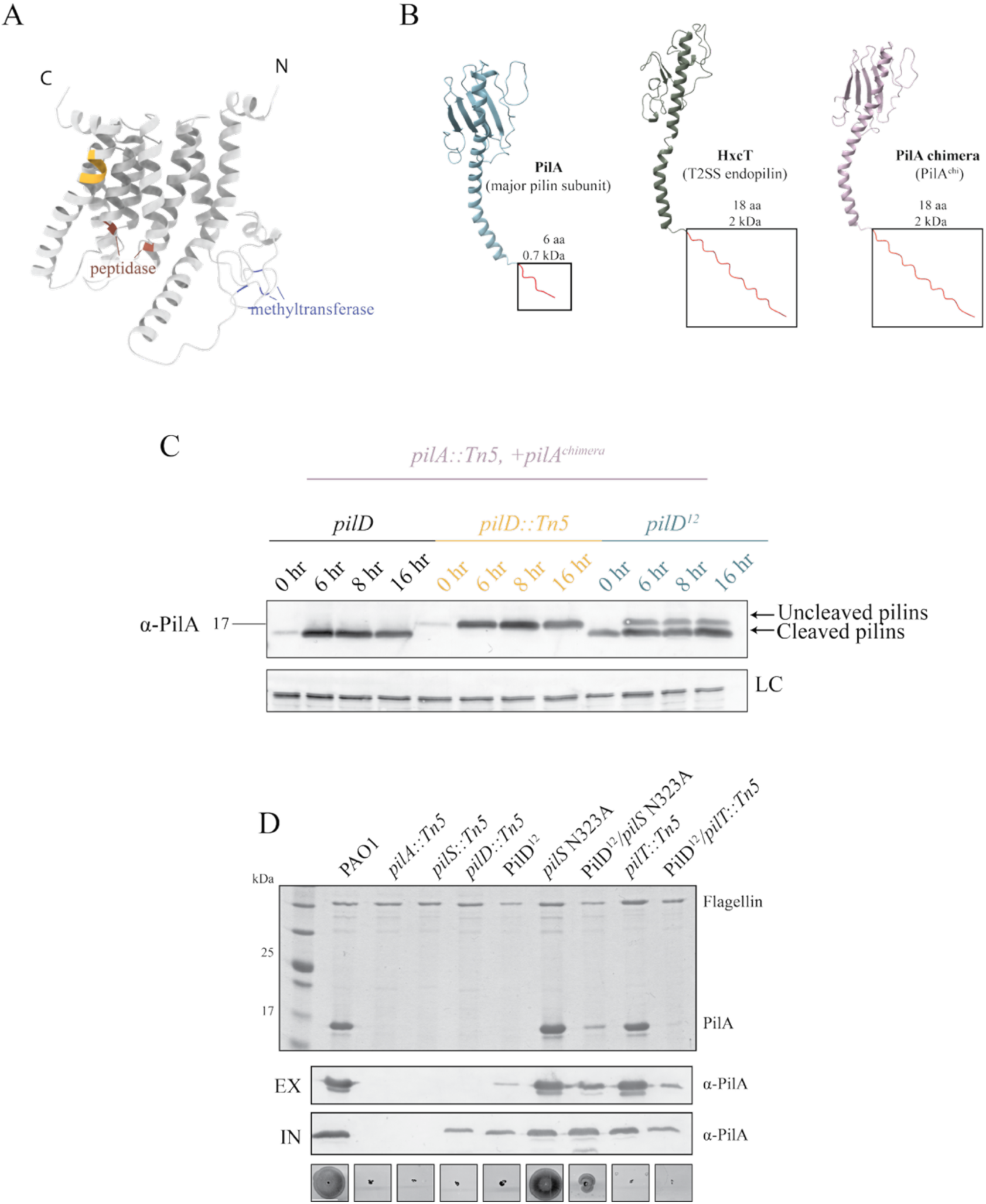
PilD^12^ mutants have reduced T4P expression and function. **A)** AlphaFold3 PilD^12^ structural model (confidence data available in **Supplementary Figure S3)**, with the 4-amino acid insertion highlighted in yellow. Key peptidase active site residues (D149, D217) are shown in blue, predicted methyltransferase residues (C72, C75, C97, C100) in purple (UCSF Chimera X). **B)** Models of PilA and the endopilin HxcT, with the leader sequences highlighted in red. Structures were predicted using AlphaFold3 (confidence data in **Supplementary Figure S2)** and modeled using UCSF Chimera X. The lengths and molecular weight of the leader sequences are indicated. The leader sequence of PilA was replaced with the HxcT leader sequence to create PilA^chimera^. **C)** In *pilD^12^*, intracellular pilins are a mixed population of cleaved and uncleaved at 6- and 8-hr post subculture, while only cleaved pilins were detected in wild type. Pilins at 16 hr were mostly cleaved. Samples were induced using 0.2% L-arabinose and separated on 16% Tris-Tricine gels followed by Western immunoblot analysis using α-PilA antisera. Blot is representative of three independent experiments. **D)** *pilD*^12^ has fewer recoverable surface pili than wild type, but introduction of the feedback-insensitive PilS N323A allele increases recoverable surface pili. Sheared surface protein samples were separated on 15% Coomassie-stained SDS-PA gel (top).

To more easily monitor the impact of this mutation on PilD peptidase function, we replaced the native 6-residue signal sequence of PilA with the 18-residue signal sequence of HxcT, the major endopilin of the Hxc T2SS in *P. aeruginosa* and also a substrate of PilD (48), creating a chimeric pilin, PilA^chimera^ (Figure 4B). Replacement with a longer signal sequence allowed for better resolution of the change in mass of the chimera following cleavage compared to wild-type PilA. Whole cell Western blot of chimeric pilins in PilD^12^ at 6- and 8-hours post subculture revealed a mixture of processed and unprocessed subunits, while the wild-type control had only processed subunits (Figure 4C). By 16-hours post subculture, all pilins were processed (Figure 4C). These data suggested that PilD^12^ cleaves prepilins less efficiently than its wild-type counterpart. Since PilD also processes the endopilins of the T2SS, we assessed protease secretion of PilD^12^. The mutants had no visible zone of clearing on skim milk plates even after 24 hours of incubation, suggesting impaired T2SS activity and supporting the hypothesis that PilD function was impaired (**Supplementary Figure S5**).

Western immunoblot using α-PilA antisera of cell lysates (bottom) shows the levels of intracellular pilins relative to controls. Data are representative of three independent experiments. Twitching motility of PilD^12^ and mutants correlate with the amount of recoverable pili. Scale bar indicates 1cm. IN=intracellular, EX= extracellular, EV= empty vector, pB= pBADGr.

Unprocessed pilins cannot be assembled, so they accumulate in the cytoplasmic membrane where they act as regulatory ligands to suppress the further expression of PilA via their interaction with the sensor kinase, PilS (39). Thus, the PilD^12^ mutation has the potential to reduce pilin availability in two ways: slowing pilin maturation and inhibiting expression of new prepilins due to the resulting backlog of unprocessed subunits. To test if this potential bottleneck could be alleviated by increasing the amount of prepilins produced, we introduced a PilS N323A point mutation into the *pilD^12^*background. PilS N323A lacks the ability to dephosphorylate the response regulator PilR when PilA accumulates, allowing *pilA* expression to continue (39). PilS N323A/*pilD^12^* had fewer recoverable extracellular pilins and less twitching motility than WT (Figure 4D), but more surface pili than the control strain, retraction-deficient PAO1 *pilT::Tn5/pilD^12^*(Figure 4D). In *pilT* backgrounds, surface piliation is maximal since any pili that can be assembled are trapped on the surface due to loss of the retraction ATPase. Low surface pili levels in PAO1 *pilT::Tn5/pilD* suggests that there is a pilus assembly defect in PilD^12^ strains that can be partially rescued through PilR hyperactivity.

Because introduction of PilS N323A restored twitching motility, we next evaluated if phage susceptibility was also rescued. PilS N323A is predicted to increase pilin production, so as an additional control we tried expressing *pilA* from an arabinose-inducible promoter in the *pilD^12^* background. PO4 susceptibility was assessed in a co-culture assay, where phages and bacteria were mixed and spotted onto agar. PilS N323A/*pilD^12^* was susceptible to PO4 killing, but unexpectedly, *pilA* overexpression *in trans* in *pilD^12^*did not restore PO4 susceptibility (Figure 5A). Similarly, in a standard plaque assay, PilS N323A/*pilD^12^* was susceptible to PO4 and two other T4P-targeting phages while PilD^12^ + pBADGR-*pilA* was resistant (Figure 5B).

**Figure 5.**
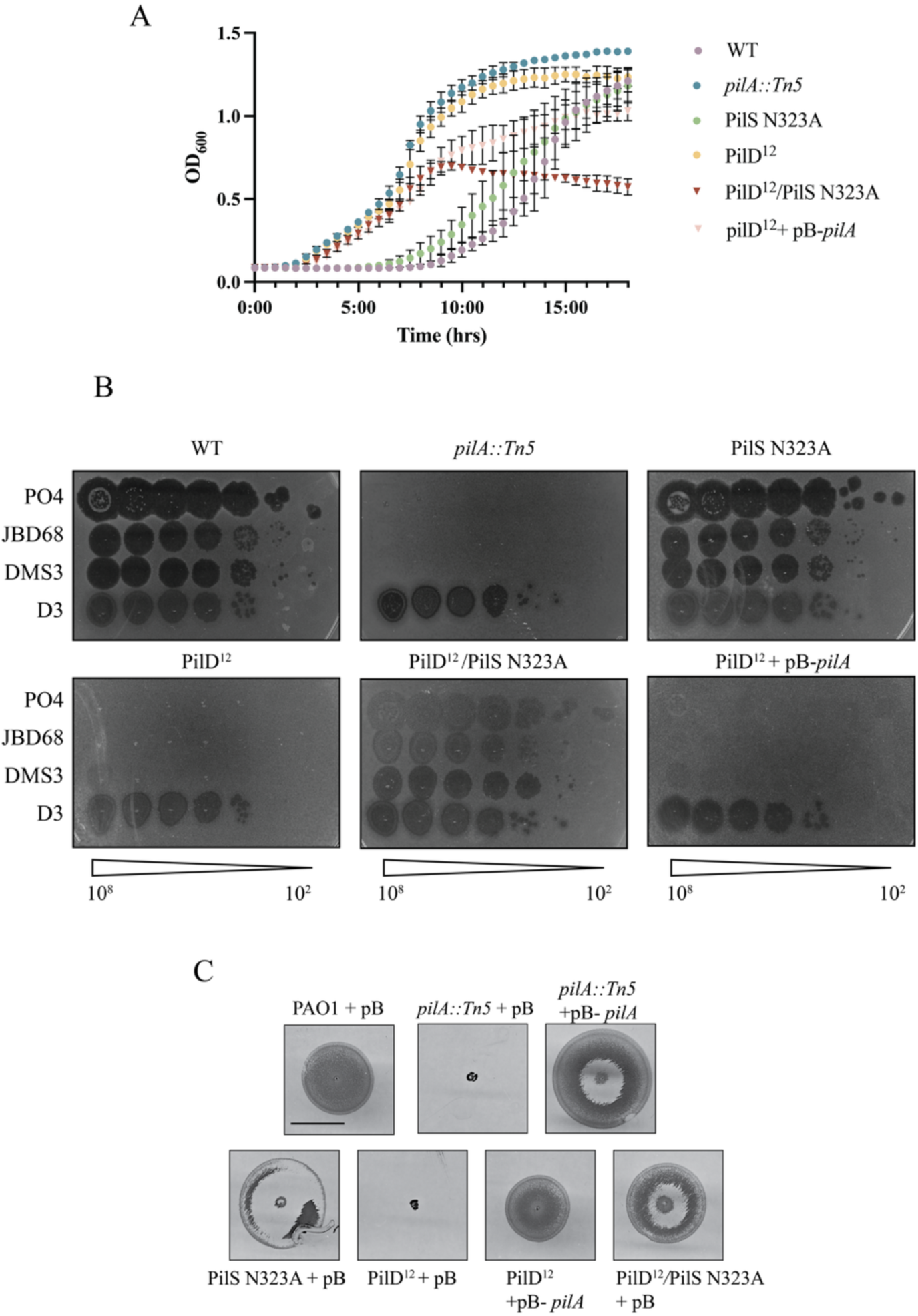
PilS N323A restores both twitching motility and phage susceptibility in PilD^12^ strains while overexpression of *pilA in trans* restores only motility. A) PilD^12^/PilS N323A is susceptible to PO4 while PilD^12^ + pBADGR-*pilA* is resistant in a liquid co-culture phage killing assay **B)** PilD^12^/PilS N323A is susceptible to 2 other T4P targeting phages while PilD^12^+ pBADGR-*pilA* is partly susceptible only to DMS3. D3 is a LPS-specific phage control. Phage titres (PFU) are indicated below each set of plaque assays. C) Twitching motility, but not PO4 susceptibility, was recovered by overexpression of *pilA in trans* in PAO1 *pilD^12^.* Samples were induced using 0.2% L-arabinose. Scale bar indicates 1 cm. Images are representative of three independent experiments. pB= pBADGr.

Interestingly, a faint plaque was observed for both PilD^12^ and PilD^12^ + pBADGR-*pilA* strains at the highest DMS3 titre tested, showing that both strains may be modestly susceptible to that phage (Figure 5B). Introduction of PilS N323A or overexpression of *pilA* from an arabinose-inducible promoter increased twitching motility in the *pilD^12^*background, although not to wild type levels (Figure 5C). These data suggest that the lack of twitching motility and decreased surface piliation in PilD^12^ is due to a combination of decreased pilin processing and negative feedback from accumulation of unprocessed pilins in the inner membrane. Both overexpression of *pilA* and PilR hyperactivity increased prepilin production and restored twitching motility in PilD^12^ but for reasons that are not yet clear, these strategies had different effects on phage susceptibility.

To determine if changing the size of the PilD insertion impacted function, we created three additional mutants with one, two, or three amino acids inserted (PilD^3^, PilD^6^, and PilD^9^) (Figure 6A). PilD^3^ twitched ∼85% (p=0.008) of WT, while PilD^6^ and PilD^9^ lacked twitching motility (Figure 6B). Intracellular pilins from PilD^3^ were processed, while those from PilD^6^ and PilD^9^ were unprocessed at all timepoints (Figure 6C), consistent with their non-twitching phenotypes. PilD^3^ phage susceptibility was comparable to that of wild-type PAO1, while PilD^6^ and PilD^9^ were resistant to all three phages tested (Figure 6D). T2SS activity was maintained in PilD^3^ but not PilD^6^ and PilD^9^ (Figure 6E). Therefore, only one or four residues can be inserted at V187 in PilD while maintaining at least partial function.

**Figure 6.**
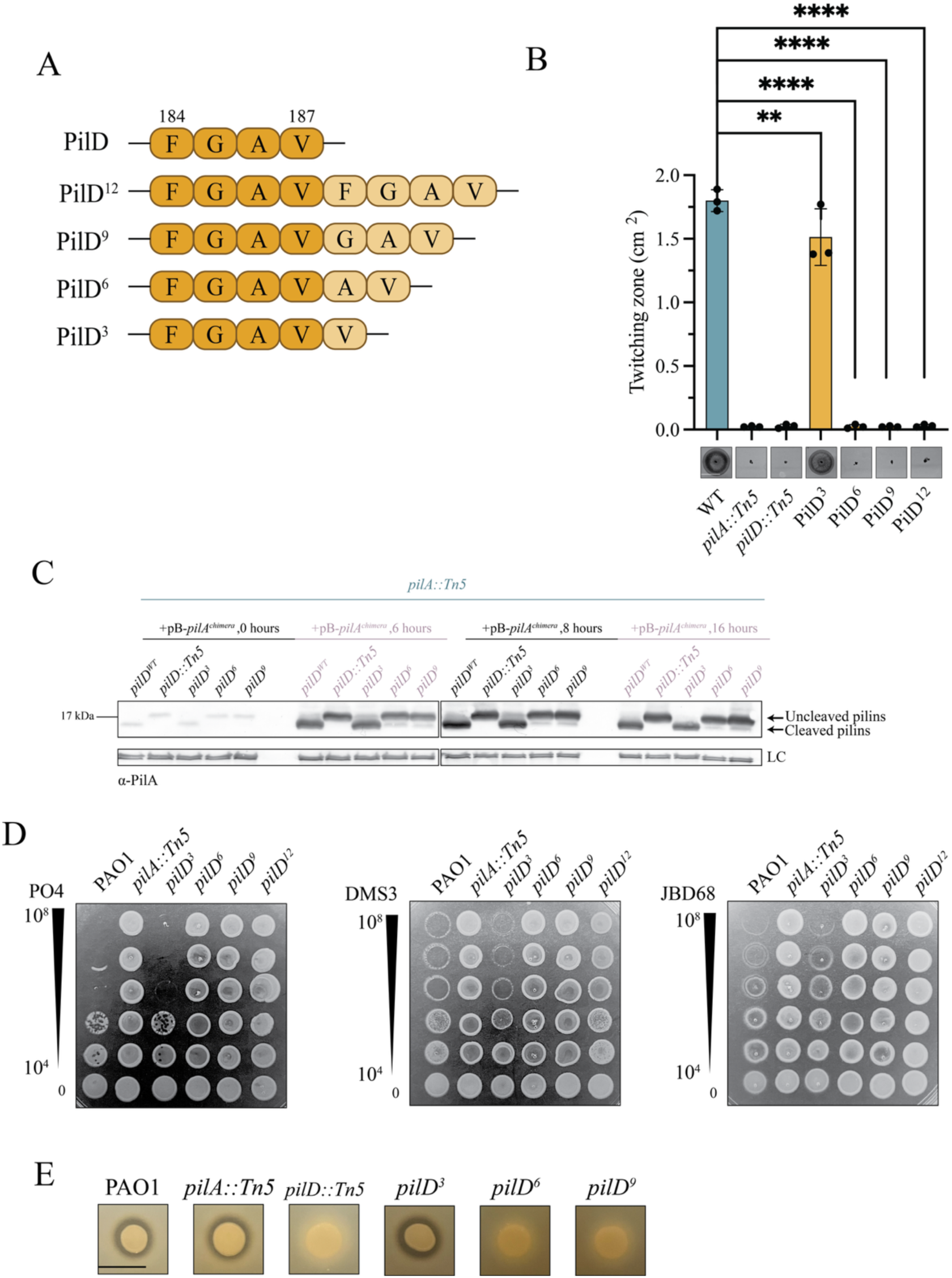
A two- or three-residue insertion at PilD V187 disrupts pilin processing. **A)** Schematic of PilD mutants. Lighter coloured residues are additions relative to wild-type residues. **B)** PilD^3^ twitches while other strains do not. Twitching zone measurements are an average of three independent experiments containing three replicates each. Twitching images are representative of three independent experiments. Scalebar represents 1cm. ** p=0.008, ****p<0.001. **C)** PilD^6^ and PilD^9^ peptidase activity is impaired as intracellular pilins were uncleaved at all time points, while PilD^3^ is functional. Samples were loaded on 17% Tris-Tricine gels followed by Western immunoblot analysis using α-PilA antisera. Flagellin levels were used as loading control. Blot is representative of three independent experiments. Samples were induced using 0.2% L-arabinose. **D)** PilD^3^ is susceptible to T4P-targeting phages while PilD^6^ and PilD^9^ are resistant. Bacteriophage plaque assays were performed using serially diluted stocks of phages PO4, DMS3, and JBD68. Titres (PFU) are indicated to the left. Image is representative of three independent experiments. **E)** *pilD^3^* spotted on a skim-milk agar plate had a visible zone of clearance, indicating protease secretion. No zone is visible in *pilD^6^* and *pilD^9^*suggesting impaired T2SS function. Scale bar represents 1cm. Samples are representative of three independent experiments.

### Only mutants with sequence duplications readily regain pilus function

For clinical applications, it is important that phage-resistant strains do not regain pathogenicity after treatment has ended, as this could contribute to recurrence of infection. Therefore, we evaluated if the PRMs recovered here could regain twitching motility, a key virulence trait, in the absence of phage pressure. Four representatives of the types of mutations that were commonly recovered in our screen were randomly selected for analysis. These included *pilT* Δ568-571 (deletion), *pilQ* C1282T (nonsense mutation), *pilB* A832C (point mutation), and PilD^12^ (duplication). Each of the mutants were first tagged with a gentamicin resistance gene as described in the Methods to ensure that any twitching colonies recovered were descendants of the relevant strains. Ten colonies of each mutant were stab-inoculated into standard 1% LB twitching agar plates. After a week of incubation at room temperature, only PilD^12^ colonies yielded twitching revertants (Figure 7A). When repeated with larger numbers, almost half of PilD^12^ colonies (45%, n=100) regained twitching motility. Since second-site suppressor mutations are more likely to occur in bacteria versus reversion to wild-type sequences (49), the *pilD* gene was sequenced. All sequenced twitching colonies had wild-type *pilD* sequences. As a control, we generated a PilD^12^ variant (PilD^12Var^) with the same 4 amino acids inserted but using different codons (Figure 7B). None of the PilD^12Var^ colonies tested regained twitching motility following a similar week-long incubation (Figure 7C). These data suggest that reversion in *pilD^12^* was likely facilitated by slipped strand mispairing that resolved the original duplication event (50).

**Figure 7.**
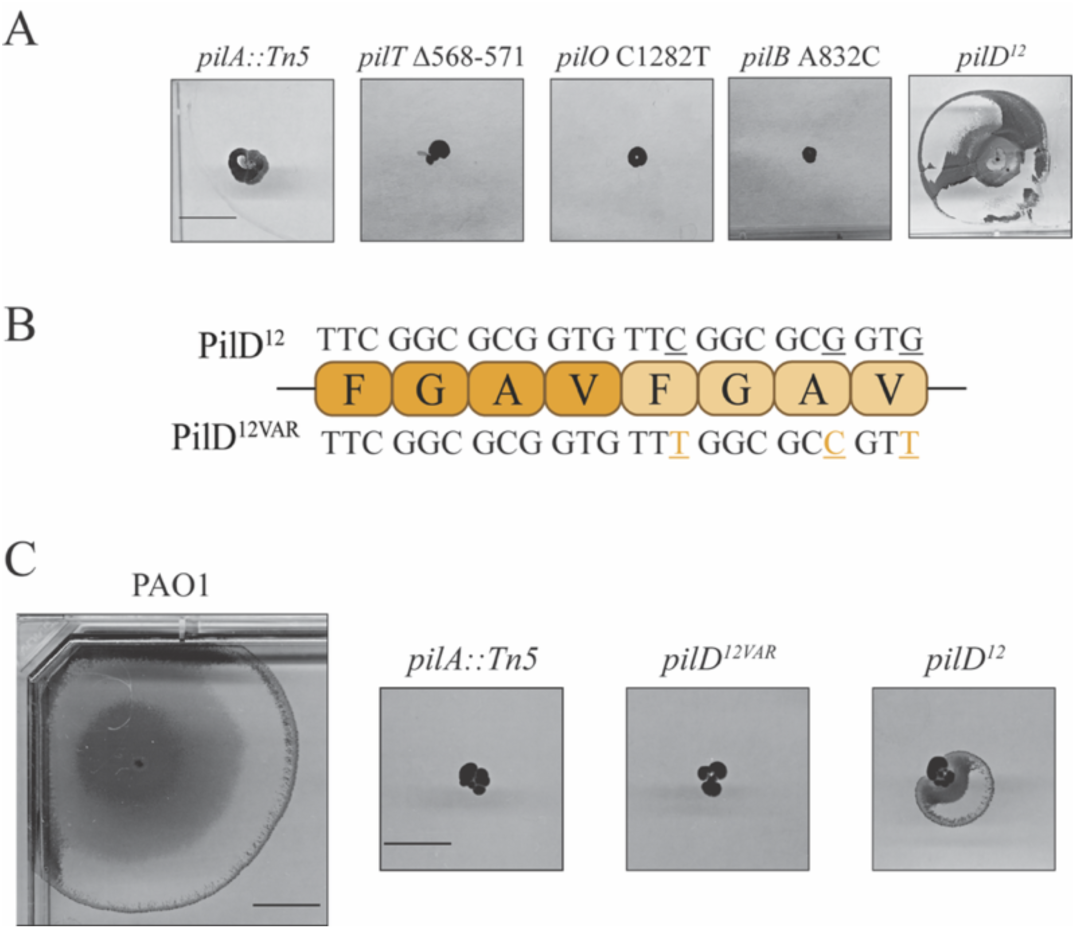
Only strains with sequence duplications readily regain motility. **A)** Randomly selected PRMs representative of each form of mutation isolated in our initial screen underwent a seven-day twitching assay, and only PilD^12^ regained motility. **B)** Schematic of PilD ^12VAR^ mutant. Lighter residues are additions relative to wild-type residues (in dark). Bases are indicated above and below amino acids with altered bases underlined in orange. **C)** PilD^12^ and PilD^12VAR^ underwent a seven-day twitching assay. Images are representative of three independent experiments. Scale bar represents 1 cm.

### A subset of resistant non-twitching strains had no identifiable mutations

Of the 128 individual PRM colonies selected, eight initially had no detectable genomic variants supported by our defined cutoff of >50 reads, which is considered sufficient for mutation identification (51, 52). Since they were not ‘mutants’ in the traditional sense, we called these isolates ‘resistant and non-twitching strains’ (RANTS). In phenotypic analyses, three RANTS expressed no pilins, three had no recoverable surface pili but produced pilins intracellularly, and two expressed surface pili (**Supplementary Figure S2)**. These data suggested that the RANTS were not clonal. All RANTS were resistant to the other T4aP-targeting phages, DMS3 and JBD68 (Figure 8A).

**Figure 8.**
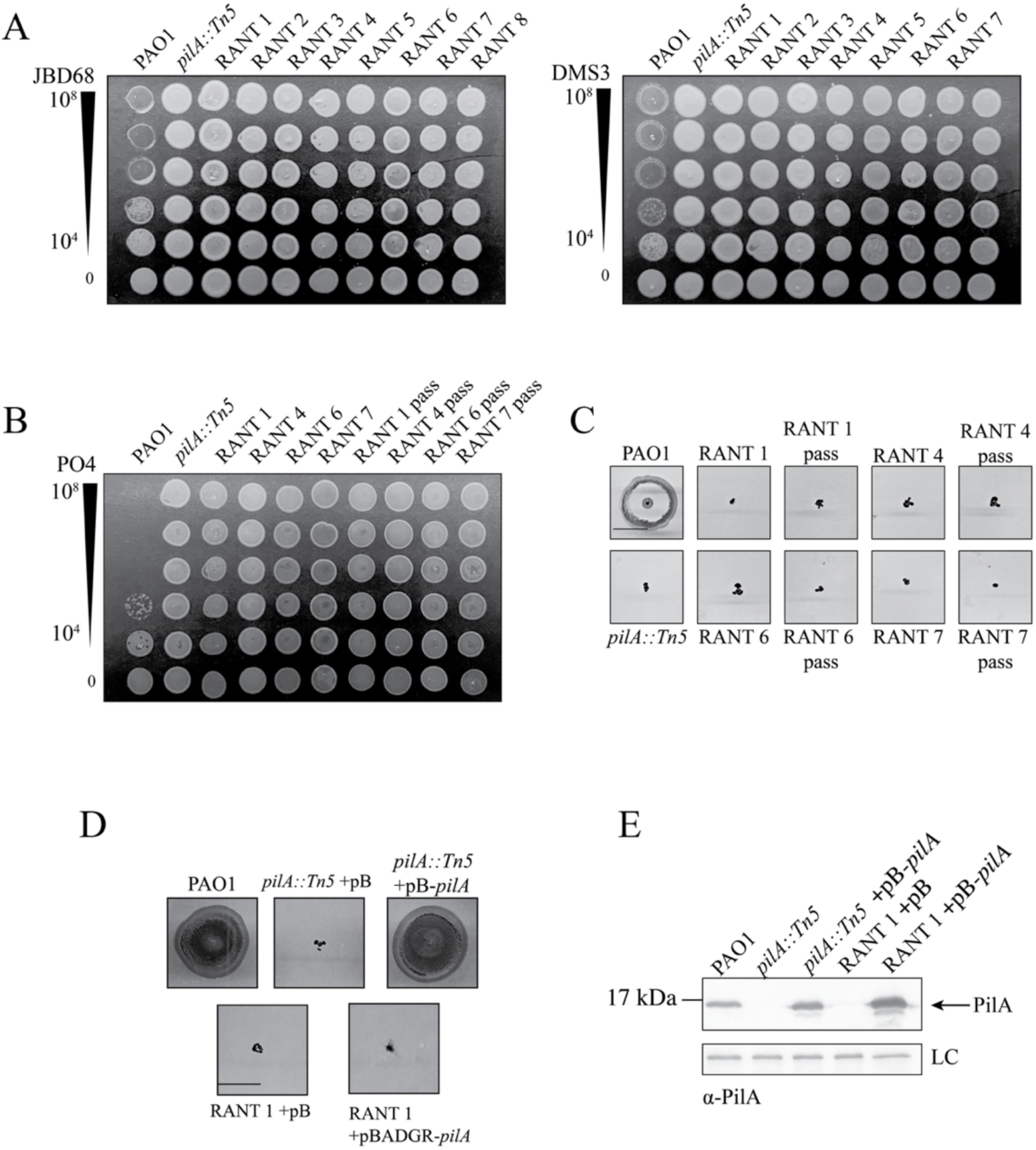
RANTS are resistant to other T4P-targeting phages and phenotypes are stable following passage in phage-free media. **A)** All RANTS are also resistant to T4P-target phages DMS3 and JBD68. Bacteriophage plaque assays using serially diluted stocks of phages PO4, DMS3, and JBD68. Titres (PFU) are indicated to the left. **B)** Following 14-day passaging in phage free nutrient-rich media, passaged RANTs (denoted with “pass”) maintained PO4 resistance and **C)** lack of twitching motility. **D)** Overexpression of *pilA in trans* in RANT 1 failed to restore twitching motility, suggesting that it has additional defects besides lack of pilin expression. Scale bar represents 1cm. **E)** Western immunoblot analysis of *pilA* expression using α-PilA antisera was used to confirm *in trans* expression. Samples were induced using 0.2% L-arabinose. Samples are representative of three independent experiments.

To rule out technical errors, three randomly selected RANTS were re-sequenced and analyzed, plus those second-round sequences were concatenated with the first-round sequences to increase the read depth. Despite this increased sequence coverage, we still could identify no mutations above our cutoffs. While breseq uses a conservative, binary approach to identify mutations such as SNPs and indels, it completes a secondary analysis that considers intermediate mutations, those that do not pass the binary cutoff but where the sequence varies from the reference, to account for mixed genomic populations or potential sequencing issues (53–55). Mutations identified using this secondary form of analysis are classified as “marginal” mutations. Using this secondary analysis, relevant mutations could be mapped to four of the eight RANTS (**Table 2**).

**Table 2.** Secondary RANT BreSeq analysis results.

| **RANT** | **Phenotype (EX/IN)<sup>a</sup>** | **Gene mutated<sup>b</sup>** | **Effect (gene)** | **Location of mutation (Total AA length)** | **Effect on protein** |
| --- | --- | --- | --- | --- | --- |
| RANT 1 | X/X | N/A |  |  |  |
| RANT 2 | X/X | *pilR* | (+1018A) | Q40FS | longer transcript |
| RANT 3 | X/ YES | *pilB* | ΔT646 | F216FS | Frameshift-shortening |
| RANT 4 | X/ YES | N/A |  |  |  |
| RANT 5 | X/X | *pilR* | T415C | PS139P | point mutation |
| RANT 6 | YES/YES | N/A |  |  |  |
| RANT 7 | X/YES | N/A |  |  |  |
| RANT 8 | YES/YES | *pilR* | A823C | T275P | point mutation |

<sup>a</sup> EX=extracellular pili, IN=intracellular pilins, X=none detected, YES=detected on gel/blot
<sup>b</sup> N/A=no change in genomic sequence compared to the wild-type reference

To assess the stability of resistance in the remaining RANTs, they were passaged daily in phage-free LB media. After 14 days of passage, all remained PO4-resistant and non-twitching (Figure 8B**, 8C)**, suggesting that the phenotypes are stable in the absence of phage pressure. As a final confirmation of sequencing quality, *pilA, pilS,* and *pilR* were sequenced in the RANTs that did not produce pilins, as these genes contribute to PilA production. All had wild-type sequences, confirming that mutations in those genes were not responsible for phage resistance. Finally, we tested whether the resistant phenotype of RANT1, which did not produce detectable pilins, could be rescued by *pilA* expressed *in trans*. Neither pilus function nor phage susceptibility was restored in the RANT1 background (Figure 8D) even though pilins were successfully expressed from the plasmid (Figure 8E), suggesting that both pilin expression and pilus assembly were perturbed in that strain.

## DISCUSSION

LPS and T4P are the most common phage receptors for *P. aeruginosa* (56–58), and most therapeutic cocktails include phages that recognize those surface structures. Here, we evaluated the repertoire and stability of mutations conferring resistance to a lytic T4P-targeting phage, PO4. This phiKZ-like phage has been used as a reporter of *P. aeruginosa* pilus function for decades and was originally used to identify putative genes involved in T4P biogenesis (18, 25, 59). Although initially reported to be specific for the PAK strain (7), our stock can infect PAO1 as well as PAK. We found mutations conferring PO4 resistance in genes encoding most structural components of the T4P, plus the PilSR TCS, and all resulted in loss of twitching motility. No hypothetical or unannotated genes associated with phage resistance and/or loss of twitching motility were identified in our analysis of this well-characterized strain. With the exception of PilD^12^, the subset of mutants we tested remained stably resistant over one week of growth in the absence of phages. A limitation of this study is use of non-selective growth conditions, where loss of T4P does not impose a fitness cost. In the more complex environment of the host, selection of suppressors that maintain fitness may be more likely.

In addition to revealing how bacteria can become resistant to phages, the serendipitous mutations that are selected can provide new insights on the roles of pilus machinery components and the residues that are key for their function. It seems unlikely that there are genetic “hot spots” for mutations that confer resistance to T4P-targeting phages given the distribution of mutations we found in a variety of genes. Genomic factors such as operon order, role of the protein, size and sequence of the gene, and additional factors such as effects on fitness may contribute to likelihood of mutation selection. Although we looked for them, we found no phage-resistant mutants that retained the ability to twitch, suggesting that loss of pilus function or expression is by far the most likely route to resistance.

At least 40 genes are directly or indirectly involved in *P. aeruginosa* T4P function and regulation (3, 11). We identified mutations in 16 unique genes, encoding members of the T4P alignment subcomplex, PilD, the PilSR TCS, some minor pilins, and two out of three ATPases. Most of the genes identified encode structural or enzymatic components, and of those, *pilD* is the only one shared with another system, the T2SS (47). When considering which pilus genes were not found in our screen, it is important to note that mutation of only a subset of known pilus genes is expected to result in phage resistance and/or loss of twitching motility. For example, the auxiliary retraction ATPase PilU is required for twitching motility but not phage infection, showing that the phenotypes can be separated (18). Components of the regulatory Pil-Chp chemotaxis system that increases intracellular cAMP levels in response to pilus surface sensing were not identified in our screen (60). However, a mutation in the Pil-Chp chemoreceptor PilJ was reported in PRMs resistant to another T4P-targeting phage (61). *pilJ* mutants have low levels of cAMP and thus reduced expression of surface pili, which could impact their susceptibility to phages that are sensitive to receptor abundance (62, 63). Most mutations we identified were SNPs or small indels, though some large genomic deletions were also identified. These patterns are comparable to those in other studies that reported mutations in *P. aeruginosa* PRMs (37, 61, 64, 65).

Three PRMs identified here had mutations in *pilB,* a frequently reported hit in other studies of phage-resistant mutants (37, 61, 64, 65). Interestingly, one of the PilB mutations identified here, PilB T278P, was also reported in PRMs isolated by another group, using a different phage (65). The other PilB mutation we identified, PilB D388A, is adjacent to the Walker B motif and the mutant produced reduced levels of surface pili that did not support motility (Figure 2). While D388 is not directly involved in Mg^2+^ coordination, this mutation may lead to structural changes in PilB that indirectly affects its ATPase activity, reducing pilus assembly. A previously described point mutation (E354Q) in a conserved residue of the Asp Box involved in Mg^2+^ coordination, also important for ATPase activity, resulted in reduced twitching motility (66). Phage susceptibility of this mutant was not assessed. These examples suggest that mutations that reduce but do not abolish PilB function are sufficient to decrease pilus assembly, potentially leading to phage resistance. It is also notable that PilB and its regulatory partners such as PilZ are favoured targets of various prophage-encoded inhibitory proteins that impair pilus function to provide super-infection exclusion (67, 68). PilB may be a common prophage target because pilus extension is energetically costly, with each pilin incorporation event requiring two molecules of ATP (40). Therefore, inhibition of PilB activity may improve growth and fitness of the lysogen. Mutations in *pilB* have been identified in clinical isolates, both independent of and following phage therapy (69, 70). Although the specific mutations are not always defined, these data suggests that such strains can survive *in vivo*.

We also recovered a PRM with a 12-base duplication in *pilD*. We showed that PilD^12^ has impaired pilin processing, with mixed populations of mature and uncleaved pilins at 6- and 8-hours post induction. The PilD^12^ uninduced overnight samples (0 hr) (Figure 4D) had more pilins than the WT *pilD* and *pilD::Tn5* uninduced overnight samples. This reproducible phenotype **(Supplementary Figure S4)** suggests that PilD^12^ may increase pilin inner-membrane levels. We hypothesize that the reduced processing capacity of PilD^12^ limits the number of mature pilins available for incorporation into the growing pilus and thus fewer surface-exposed pili. Simultaneously, the prepilins that accumulate repress further PilA expression via their interactions with the PilSR system (Figure 9A). Together, these events lead to a pronounced decrease in availability of mature pilins for pilus incorporation. Combining a PilS N323A mutation with PilD^12^ addressed these issues by increasing the overall amount of prepilins (Figure 9B), restoring motility and phage susceptibility (Figure 4D). However, expressing PilA from a plasmid, which also increases overall amount of prepillins, restored twitching motility but not phage susceptibility (Figure 5). In PilS N323A, PilR is locked in its phosphorylated, active state; in contrast, overexpression of PilA from a plasmid leads to PilR dephosphorylation by PilS. Our data suggest that increasing pilin abundance in PilD^12^ is sufficient to restore twitching motility, but successful phage replication may also require PilR activation. In addition to *pilA*, PilR regulates a large set of genes including those encoding the minor pilin subcomplex and PilBCD (71). Increasing pilin production through the PilSR TCS coordinately upregulates other pilus genes to produce more pili that phages can recognize. Phage replication is also affected by cell metabolism and growth (72, 73). PilR also regulates a subset of host metabolic enzymes that may similarly contribute to phage replication (71).

**Figure 9.**
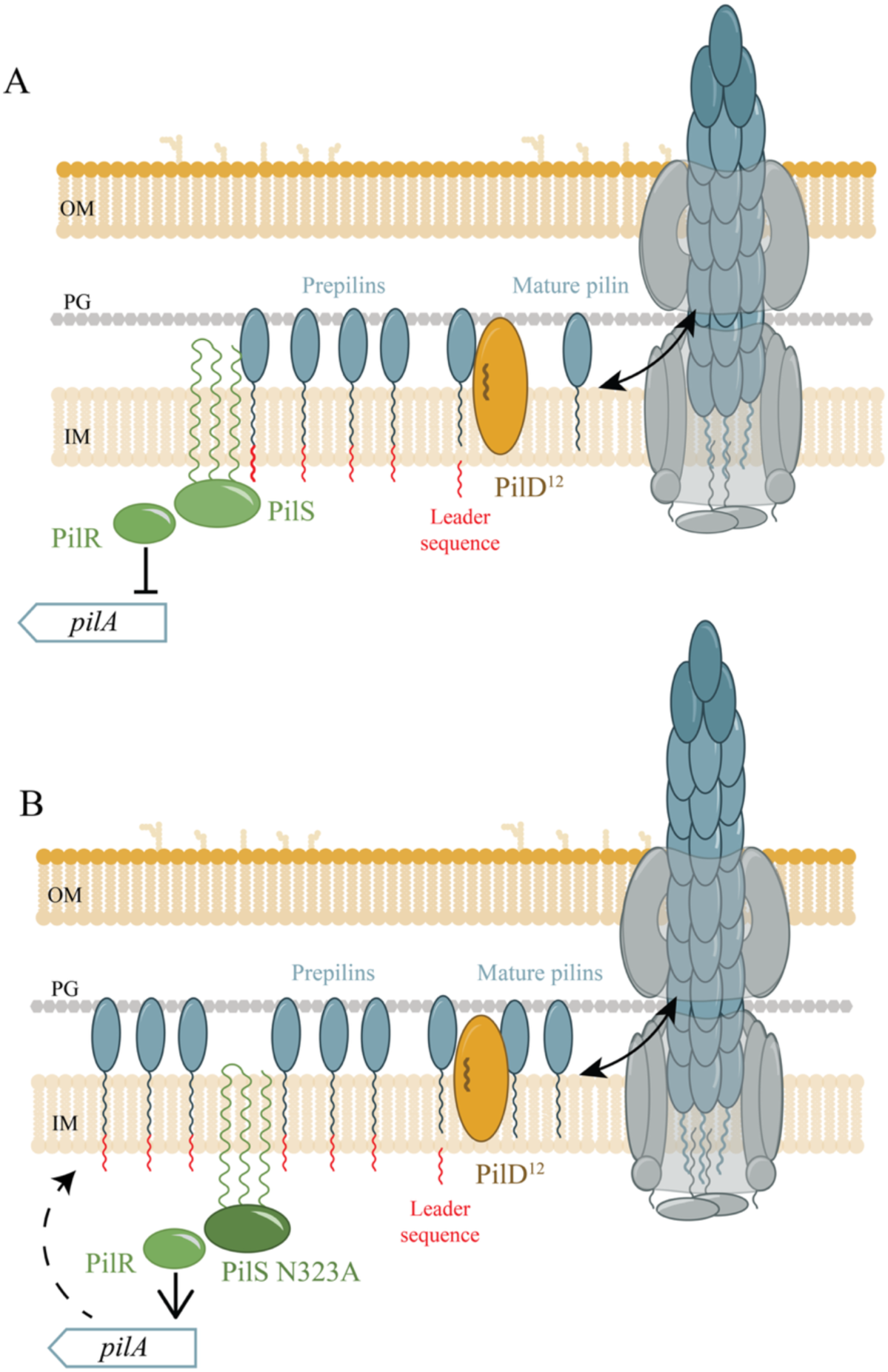
Model of PilS-dependent alleviation of the PilD^12^ decrease in pilus assembly. **A)** In PilD^12^ background, accumulation of prepilins in the inner membrane is sensed by PilS which dephosphorylates PilR, decreasing *pilA* expression. This results in production of fewer pre-pilins, further reducing the pool of mature pilins that can be assembled into pili. **B)** In PilS N323A, feedback inhibition by prepilins is lost and phospho-PilR continues to promote *pilA* expression, increasing the total pool of prepilins. This allows for more pilins to be processed by PilD^12^, increasing pilus assembly.

Based on structural predictions, the PilD^12^ duplication is located in the fifth transmembrane helix of PilD, distal from the cytoplasm-facing peptidase and methyltransferase active sites (Figure 4A). One complete turn of an α-helix requires ∼4 amino acids to maintain correct hydrogen bonding (74). Inserting two or three residues in PilD^6^ and PilD^9^, respectively, likely disrupts hydrogen bonding, affecting protein stability or its interaction with pilins. Those two mutants resembled a strain lacking PilD in terms of phage susceptibility, motility, and secretion. In contrast, PilD^12^ is predicted to lengthen the helix by one complete turn, allowing the enzyme to retain partial function. Despite its essential role in function of the T4P and T2SS systems and contributions to *P. aeruginosa* virulence, PilD has been poorly characterized compared to other components (75). This interesting mutation provides new insight into allosteric control of PilD’s peptidase function.

We were surprised to isolate some strains that were stably phage-resistant and non-twitching (RANTs) where we could identify no mutations. Some were ultimately resolved by further scrutiny of mutations that did not pass the initial sequence quality and read depth cutoffs, but there were still 4 RANTS with genomes that appeared identical to their WT parent strain.

The ability of *P. aeruginosa* to become phage resistant in the absence of obvious mutations has been reported previously (76, 77). It is possible that epigenetic changes following phage exposure could result in silencing of key genes (i.e. pilus structural components, genes that directly or indirectly regulate T4P function, or combinations thereof). In *Ralstonia pseudosolanacaerum*, methylation patterns that arose during growth in the plant host persisted for up to 22 passages outside of the host, demonstrating the longevity of prokaryotic epigenetic modifications (78). Such changes might be detected using newer forms of whole genome sequencing such as PacBio or Oxford Nanopore sequencing (79, 80). Thus, future work using these more advanced sequencing technologies may shed light on how phage resistance can be achieved in the absence of detectable mutations.

The inexorable rise of antibiotic resistance has led clinicians to reconsider phage therapy as an option (81). Most published case studies show that it is successful and safe, supporting this approach (82, 83). Our findings can inform development of phage therapy and phage steering strategies. One caveat of this work is that PRMs were selected in rich media, and more host-like conditions (eg. nutrient limitation, immune system components) might affect the types of mutations that are selected. However, phage resistance patterns identified *in vitro* appear to accurately reflect those arising *in vivo* (70).

Our data suggest that T4P-targeting phages mostly select for mutations causing the irreversible loss of a key virulence factor. Strains lacking T4P have decreased pathogenicity in different *in vivo* models (58, 84–86), therefore this phenotype may be clinically beneficial.

Ideally, T4P-targeting phages should be combined with phages using other receptors, such as LPS, to address resistance and emergence of potential revertants. This strategy has been applied in phage cocktails BFC1 and 2, created by the Queen Astrid Military Hospital in Belgium. Both contain T4P-targeting phage PNM, along with a LPS-targeting phage and other phage(s) with unknown targets (87–89). These cocktails as well as PNM alone have been successful in clearance of infection (83).

T4P are common surface structures that are expressed by other important human pathogens, including *Acinetobacter baumannii*, *Stenotrophomonas maltophilia, Neisseria meningitidis, Streptococcus pneumoniae,* and *Clostridioides difficile* (2, 90–92). Pilus-targeting phages for many of these pathogens have been identified (8, 93). Similarly, plant pathogens *Xylella fastidiosa, Xanthomonas axonopodis, Xa. euvesicatoria,* and *P. syringae* express T4P, and pilus-targeting phages have been identified for those species as well (94, 95). The use of phages in agriculture for biocontrol is a growing field (96, 97). It will be interesting to learn whether T4P-targeting phages for other host species will select for similar repertoires of escape mutations.

## METHODS

### Bacterial strains and growth conditions

Bacterial strains were grown in lysogeny broth (LB) at 37°C with shaking (200 rpm) or on 1.5% agar plates supplemented with antibiotics with the following concentrations, when required: gentamicin (Gm) 30 μg/mL, carbenicillin (Carb) 200 μg/mL, or gentamicin (Gm) 15 μg/mL for *E. coli*. L-arabinose was added to media at a concentration of 0.2% (w/v), when required, to induce expression from the pBADGr Ara promoter. All bacterial strains and plasmids used for this study are listed in **Table S1.**

### Phage genome sequencing and analysis

Bacteriophage genomes were extracted and assembled as previously described (62). Phage PO4 was amplified using *P. aeruginosa* PAO1 as the host strain. Phage genomic DNA was extracted (Norgen Biotek Phage DNA Isolation Kit) and sequenced using the Illumina NextSeq 2000 platform (Microbial Genome Sequencing Center/SeqCenter, Pittsburgh, PA, USA).

Sequencing results were assembled using SPAdes (version 3.15.5) (98) and annotated using Pharokka (version 1.7.3) (99). The annotated genome of PO4 is available at the following GenBank accession number: PX259653.

### Molecular biology

All primers used are listed in **Table S2.** Chromosomal mutants were generated as previously described (100) with the exception of pEX18Gm constructs with *pilD^3^*, *pilD^6^*, *pilD^9^*, and *pilD^12VAR^*. These *pilD* variants were synthesized (gBlock, IDT). The oligos were digested with the indicated restriction enzymes and ligated into pEX18Gm digested using the same restriction enzymes. Plasmid and chromosomal sequences were confirmed using Sanger sequencing (McMaster Genomics Facility) or Oxford Nanopore whole-plasmid sequencing (Plasmidsauras).

The PilA chimera construct was created by PCR amplification of PAO1 *pilA* using an upstream primer that contained the leader sequence of *hxcT* and first 15 bases of mature PAO1 *pilA* and downstream primer corresponding to the end of PAO1 *pilA* (Table S1). Amplified genomic DNA was digested with appropriate restriction enzymes and ligated to pBADGr digested with the same enzymes. The plasmid was introduced into PAO1 *pilA::Tn5* by electroporation as described previously (100) and transformants selected on plates containing Gm as above.

### Selection of phage-resistant mutants

*P. aeruginosa* PAO1 was inoculated from overnight subculture into 5 mL LB, and incubated at 37°C with shaking (200 rpm) to an optical density at 600 nm (OD600) of 0.6. When target OD600 was reached, 100 μL of the liquid bacterial culture was combined with 12 mL of 0.6% LB agar and 10 μL of PO4 at concentration of 10^8^ PFU in culture tubes and poured into petri dishes. Plates were incubated at 21°C for an additional 24 h or until the appearance of bacteriophage resistant colonies. Colonies were selected and grown on 1.5% LB agar, incubated at 37°C. To confirm bacteriophage resistance, phage streaking assays were completed using the phage resistant colonies. On 1.5% LB agar plates, 5 μL of PO4 at concentration of 10^8^ PFU was applied onto the surface in four evenly spread out spots. An inoculation loop was dipped into liquid culture of resistant colonies and dragged across the plate, crossing each phage sample. Plates were incubated overnight at 37°C and checked the following day for growth.

### Twitching motility assays

Twitching motility assays were completed as previously described (39). In brief, bacterial strains were stab-inoculated through cell-culture treated single well plates containing 1% LB agar. Standard twitching assay plates incubated at 37°C, overnight. LB was supplemented with L-arabinose and Gm when required. To visualize the twitching zone, agar was removed, and plates were stained with 1% crystal violet for 5 min before washing with water to remove excess dye. Plates were imaged using a flatbed scanner.

Samples for extended-length twitching assays used to assess phenotypic reversion were also stab inoculated through cell-culture treated single well plates containing 1% LB agar. These plates were sealed with Parafilm and placed in a sealed zipper storage bag. After seven days of room temperature incubation, in the dark, for any twitching zones present, approximately 1cm of cell growth were collected using a sterile cotton swab and transferred to Gm supplemented media to assess growth. Remaining intact twitching zones were visualized as above.

Twitching zones were measured using Image J, when indicated. Average areas were graphed using GraphPad Prism (version 10.2.0). One-way ANOVA statistical analysis was performed on the areas using GraphPad Prism (p< 0.05 is considered statistically significant).

### Whole genome sequencing and mutation analysis of PRMs

Genomic DNA of *P. aeruginosa* PAO1 WT and phage resistant strains were extracted using the Wizard Genomic DNA Purification Kit. PRMs and PAO1 WT were sequenced using the Illumina NextSeq 2000 platform (Microbial Genome Sequencing Center/SeqCenter, Pittsburgh, PA, USA). All samples had average sequencing coverage of at least 50 reads. breseq (versions 0.35.4, 0.36.1, or 0.37.1) was used to identify mutations by comparing mutants sequences to the reference WT PAO1 Burrows lab strain (53). To ensure variant calling results were reliable, only those with more than 50 reads were included in the analysis unless specified (101). When required, mutations were confirmed by PCR and DNA sequencing (Plasmidsaurus, Oxford Nanopore).

### Multi-strain phage susceptibility assays

Overnight cultures of bacteria were grown (1:1000) in liquid LB media and incubated at 37°C with shaking (200 rpm). Subcultures were standardized to OD600 of 0.5 in LB media. Phage stocks were standardized to 10^8^ PFU/mL and 10-fold serially diluted in phage buffer (68mM NaCl, 10mM Tris-HCl pH7.5, 10 mM CaCl2, and 10mM MgSO4). Five μL each of culture and phage were combined and spotted onto 0.6% LB agar plates and dried for 5 min before incubation for 18-24 hours at 21°C. Phage susceptibility is indicated by absence of cell growth, relative to the control.

### Single strain phage plaquing assays

Overnight culture of bacteria were grown (1:100) in liquid LB media and incubated at 37°C with shaking (200 rpm). Subcultures were standardized to OD600 of 0.3 in LB media and 100 μL of diluted culture was mixed with 12 mL of 0.6% LB agar and poured into standard petri dishes. LB agar was air dried in a biosafety cabinet. Phage stocks were standardized to 10^8^ PFU/mL and serially diluted with phage buffer. Five μL of each dilution was spotted onto the plates and air dried. Plates were incubated for 18 hours at 30°C. Phage susceptibility is indicated by phage plaque clearance.

### Liquid phage susceptibility assay

Strains of interest were combined with PO4 at MOI=1 in 100 μL LB. Samples were grown shaking (orbital) at 37°C for 18 hours in the Biotek Epoch Plate readers. OD600 measurements were taken every 30 minutes. OD600 measurements were graphed using Graphpad Prism version 10.2.0. Phage susceptibility is indicated by lack of growth.

### Sheared surface protein analysis

Sheared surface protein preparations were completed as previously described.^15^ Each strain/mutant was streaked on two 1.5% LB agar plates, containing L-arabinose and Gm when required, in a grid and incubated overnight at 37°C. Cells were gently scraped using sterile coverslips and resuspended into 3 mL of 1X phosphate buffered saline (PBS) to be vortexed for 15 s, twice. Samples were aliquoted into 1.5 mL Eppendorf tubes and centrifuged at 21,000 x *g* for 30 min. Supernatant was transferred into new 1.5 mL Eppendorf tubes, 5M NaCl and 30% (w/v) polyethylene glycol (8000) were each added to concentrations of 10% final volume.

Samples were incubated on ice for 60 min before centrifugation at 21,000 x *g* for 30 min to precipitate proteins. Precipitated proteins were resuspended in 50 μL of sample loading buffer (62.5mM TRIS-HCl pH 6.8, 2.5% SDS, 0.002% bromophenol blue, 5% β-mercaptoethanol, 10% glycerol) and boiled for 10 min. Samples were cooled to room temperature before separation on 15% SDS-PAGE and visualized using Coomasie brilliant blue. Where pilin proteins could not be detected on gels, their presence was confirmed using Western blot analysis, as described below.

### Whole cell protein sample analysis

Bacterial strains were streaked on 1.5% LB agar plates and incubated overnight at 37°C. Using an inoculating loop, bacteria were resuspended into 2 mL of 1X PBS and mixed. Samples were diluted to an OD600 of 0.6 and 3mL transferred into 2 1.5 mL Eppendorf tubes before centrifugation for 1 min at 21,000 x *g*. The supernatant was discarded and precipitated cells were resuspended in 50 μL of 1X loading buffer. Samples were boiled for 10 min and cooled to room temperature before separation with 15% SDS-PAGE, in 1X Tris-glycine running buffer (diluted from 10X tris-glycine buffer containing 30.3 g tris, 144 g glycine, and 20 mL 10% SDS). Following separation, proteins were transferred to a nitrocellulose membrane for 1 h at 225mA in 1X transfer buffer (20% methanol, 100mL of 10X tris-glycine buffer without SDS, in 1L of H2O) Membranes were blocked with 5% skim milk resuspended in 1X phosphate-buffer saline (PBS) for minimum of 2 h, room temperature shaking. Primary antibodies were diluted in 1x PBS and used in appropriate dilutions (α-PilA-1:5000, α-PilB-1:2000,), and incubated with the blot at room temperature overnight. Membranes were washed 4x for 5 min with PBS followed by incubation with 1:3000 dilution (1xPBS) of goat α-rabbit alkaline phosphatase-conjugated secondary antibodies for 1 h at room temperate with shaking at 60 rpm. Membranes were washed 4 x for 5 min with 1x PBS before developing using 5-bromo-4-chloro-3-indoyl phosphate (BCIP) and nitro blue tetrazolium chloride (NBT) resuspended in alkaline phosphatase buffer (1mM Tris, 100mM NaCl, 5mM MgCl2, pH 9.5), shaking in the dark for 10 minutes. Western blots were imaged using a flatbed scanner.

### Time-course intracellular pilin processing analysis

Overnight liquid bacterial cultures were grown in LB supplemented with 0.2% L-arabinose and 30 mg/mL Gm and diluted to OD600 of 0.6 for timepoint 0 h and prepared as described above. At each timepoint, cultures were diluted to OD600 of 0.6, if required, and prepared as described above. Samples were separated using 16% Tris-Tricine gels in 1x running buffer (diluted from 10X running buffer containing 24.2g Tris base, 17.9g tricine, 1g SDS) instead of 15% SDS-PAGE gels (102). Following separation, proteins were transferred to a nitrocellulose membrane for 1 h at 225 mA in 1X transfer buffer (20% methanol, 100mL of 10X tris-glycine buffer without SDS, in 1L of H2O) Membranes were blocked with 5% skim milk resuspended in 1X phosphate-buffer saline (PBS) for minimum of 2 h, room temperature shaking. Primary antibodies were diluted in 1x PBS and used in appropriate dilutions (α-PilA-1:5000), and incubated with the blot at room temperature overnight. Membranes were washed 4x for 5 min each with PBS followed by incubation with 1:3000 dilution (1xPBS) of goat α-rabbit alkaline phosphatase-conjugated secondary antibodies for 1 h at room temperate with shaking at 60 rpm. Membranes were washed 4 x for 5 min each with 1x PBS before developing using 5-bromo-4-chloro-3-indoyl phosphate (BCIP) and nitro blue tetrazolium chloride (NBT) resuspended in alkaline phosphatase buffer (1mM Tris, 100mM NaCl, 5mM MgCl2, pH 9.5), shaking in the dark for 10 minutes. Western blots were imaged using a flatbed scanner.

### Protease secretion assay

Skim milk media was created by combining 100 mL of 1.5% (w/v) skim milk powder in deionized water and 100 mL LB supplemented with 1.5% tryptic soy agar (TSA), at room temperature. (103) Media was poured onto rectangular single well plates and solidified. Bacteria were inoculated in 5mL LB, and incubated with shaking (200 rpm) to an optical density at 600 nm (OD600) of 0.5. Five μL of standardized bacterial culture was applied onto the skim milk media and incubated at 21°C for 24 hours or until the appearance of clearance zones for control samples. Plates were imaged using a flatbed scanner and quantified using ImageJ.

### Passaging and phage susceptibility of Resistant and Non-Twitching Strains (RANTS)

Overnight cultures of RANTS were subcultured (1:000) in LB and incubated at 37°C with shaking (200 rpm) for 5-8 h. Subcultures were diluted to assess phage susceptibility, as described above. From the same overnight culture, 10 μL was diluted into 5 mL of LB and incubated with shaking (200 rpm) overnight. The overnight culture was diluted to OD600 = 0.6 and phage spot assay was repeated, daily for 14 days.

### Insertion of the Gm resistance gene

Prior to assessing the capacity of mutants to regain twitching motility following extended incubations of up to 7 days, the strains were marked with a Gm resistance gene. Genomic insertion using a mini-Tn7 vector was completed as previously described (104, 105). In short, pUC18t-mini-Tn7T-Gm and pTNS1 were introduced by electroporation, and transformants were recovered for 3.5 hours in LB. Cells were plated on 1.5% LB agar containing Gm overnight and successful insertion was confirmed by growth on supplemented media.

### Data availability

The genome sequence of phage PO4 is available at NCBI GenBank accession number: PX259653.

## Acknowledgements

We thank Ikram Qaderi for assistance in phage genome annotation, Dominique Tertigas for assistance in genome analysis, and Yaser Al Moayad for technical assistance with some experiments. This work was supported by a grant from the Canadian Institutes of Health Research to LLB (PJT-169053). LLB holds the Canada Research Chair in Microbe-Surface Interactions (CRC-2021-00103) and VNT holds a PGS-D award from the Natural Sciences and Engineering Research Council of Canada.

