## Supplementary Figs S1-S5 and Tables S1-S2 for "The repertoire of resistance mutations selected by a *Pseudomonas aeruginosa* type IV pilus-targeting lytic bacteriophage"

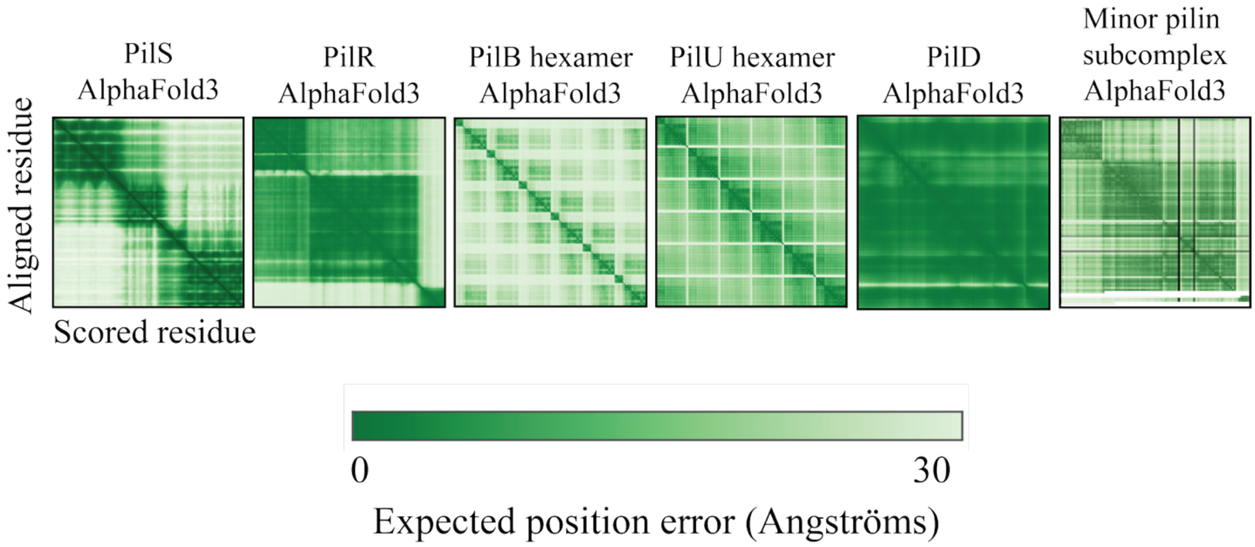


**Figure S1. Predicted aligned error plots for AlphaFold predicted structures.** Error plots for predicted structures in **Figure 1** were generated using AlphaFold3 and visualized using ChimeraX.

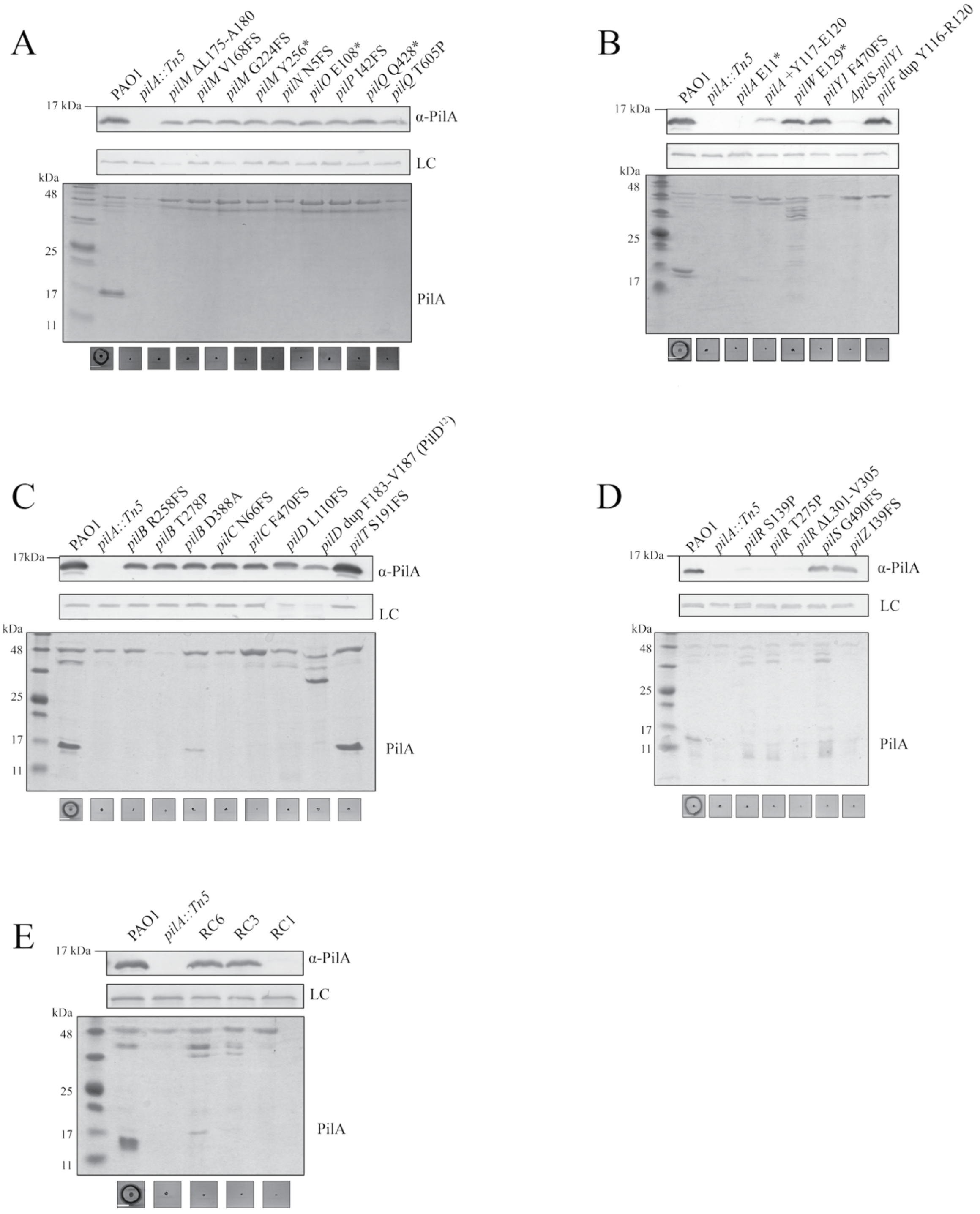


**Figure S2. PRMs do not twitch but have varied pilin production and recoverable surface pilins.** From top to bottom of each panel, Western immunoblots of cell samples representing intracellular pilin production using α-PilA antisera. Sheared surface protein preparations of respective PRMs, analyzed using a Coomasie-stained SDS-PAGE gel. Twitching analysis of each PRM, plates were incubated overnight at 37ºC. PRMs are organized based on mutations in genes of the **A)** alignment complex, **B** major and minor pilins and *pilF*, **C)** inner membrane motor complex, **D)** PilS-R TCS and PilZ, and **E)** representative RANTS. Samples are representative of three independent experiments. Scale bar represents 1cm.

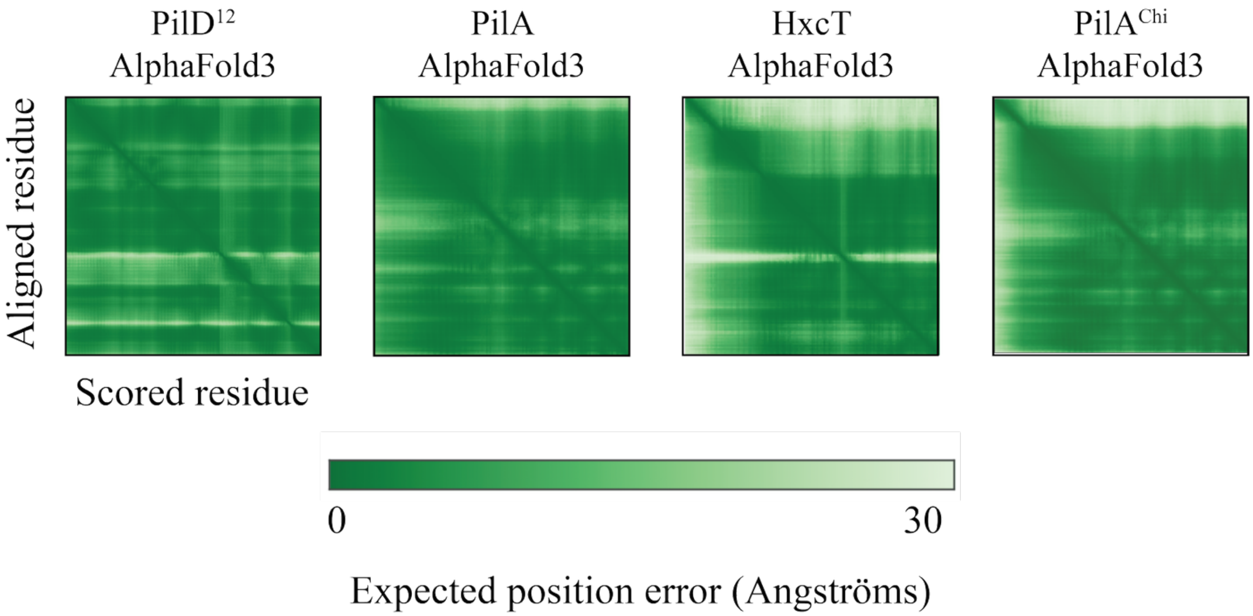


**Figure S3. Predicted aligned error plots for AlphaFold predicted structures.** Error plots for predicted structures in **Figure 3** were generated using AlphaFold3 and visualized using ChimeraX.

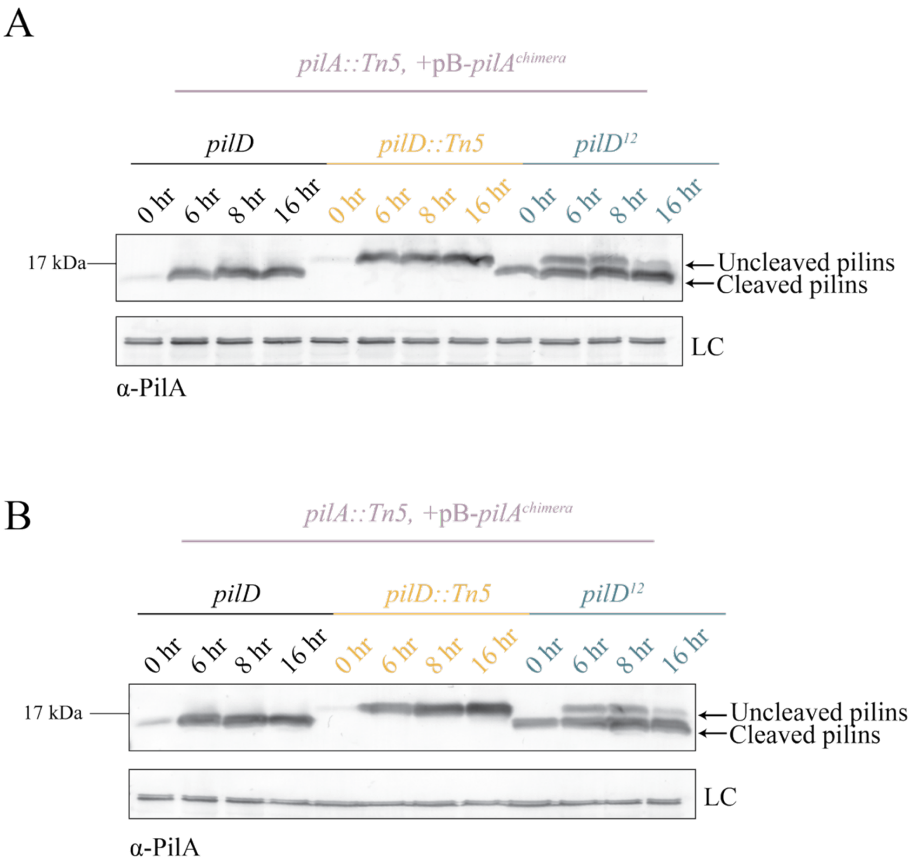


**Figure S4**. **Representative replicates of Figure 3C.** PilD$^{12}$ has more recoverable pilins than WT and NP at initial sample.

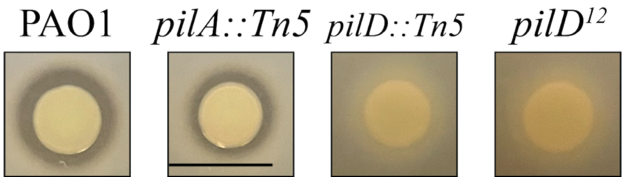


**Figure S5. PilD$^{12}$ protease secretion is impaired.** PilD$^{12}$ spotted on a skim-milk agar plate does not have a visible clearance zone, suggesting lack of T2SS-dependent secreted proteases. Scale bar represents 1cm. Samples are representative of three independent experiments.

**Supplementary Table S1. Strains and plasmids used in this study**

| Strain/plasmid | Characteristics | Source |
|---|---|---|
| *P. aeruginosa strains* | | |
| mPAO1 | WT | lab collection |
| mPAO1 + pBADGR | WT complemented with pBADGR | This work |
| mPAO1 + pBADGR-PilB D388A | WT complemented with PilB D388A/*pilB A1163C* | This work |
| mPAO1 *pilA::Tn5* | ISphoA/hah insertion at position of 163 in *pilA* | (1) |
| mPAO1 *pilA::Tn5* + pBADGR | ISphoA/hah insertion of *pilA* complemented with pBADGr | This work |
| mPAO1 *pilA::Tn5* + pBADGR- *pilA* | ISphoA/hah insertion of *pilA* complemented with *pilA* | This work |
| mPAO1 *pilB* :: Tn5 | ISphoA/hah insertion at position of 411 in *pilB* | (1) |
| mPAO1 *pilB* :: Tn5 +pBADGr | ISphoA/hah insertion in *pilB* complemented with pBADGR | This work |
| mPAO1 *pilB* :: Tn5 +pBADGr-*pilB* | ISphoA/hah insertion in *pilB* complemented with *pilB* | This work |
| mPAO1 *pilB* :: Tn5 +pBADGr-PilB D388A | ISphoA/hah insertion in *pilB* complemented with PilB D388A | This work |
| mPAO1 PilB D388A +pBADGr | mPAO1 PilB D388A/*pilB A1163C* complemented with pBADGr | This work |
| mPAO1 PilB D388A +pBADGr-*pilB* | mPAO1 PilB D388A/*pilB A1163C* complemented with *pilB* | This work |
| mPAO1 $pilD^{12}$ | 12 base duplication of residues 548-559 of *pilD* | This work |
| mPAO1 $pilD^{12}$ + pBADGR | 12 base duplication of residues 548-559 of *pilD* complemented with pBADGR | This work |
| mPAO1 $pilD^{12}$ + pBADGR | 12 base duplication of residues 548-559 of *pilD* complemented with *pilA* | This work |
| mPAO1 $pilD^{3}$ | 3 base duplication of residues 548-550 of *pilD* | This work |
| mPAO1 $pilD^{6}$ | 6 base duplication of residues 548-553 of p*ilD* | This work |
| mPAO1 $pilD^{9}$ | 9 base duplication of residues 548-556 of *pilD* | This work |
| mPAO1 $pilD^{12var}$ | Insertion of TTTGGCGCCGTT following 548 of *pilD* | This work |
| mPAO1 *pilD::Frt* | ISphoA/hah insertion at position of 416 in *pilD* | (1) |
| mPAO1 *pilD::Tn5*/*pilA* '+pBADGR'-$pilA^{chi}$ | ISphoA/hah insertion of *pilD*, deletion of *pilA,* complemented with *pilA-hxcT chimera* | This work |

| | | |
|---|---|---|
| mPAO1 *pilD*$^{3}$/*pilA* '+pBADGR'-*pilA*$^{chi}$ | 3 base duplication of residues 548-550 of *pilD*, deletion of *pilA*, complemented with *pilA-hxcT chimera* | This work |
| mPAO1 *pilD*$^{6}$/*pilA* '+pBADGR'- *pilA*$^{chi}$ | 6 base duplication of residues 548-553 of p*ilD*, pilA ISphoA/hah insertion at position 163 in *pilA*, complemented with pilA-hxcT chimera | This work |
| mPAO1 *pilD*$^{9}$/*pilA* '+pBADGR'- *pilA*$^{chi}$ | 9 base duplication of residues 548-556 of *pilD, pilA ISphoA/hah insertion at position 163 in pilA, complemented with pilA-hxcT chimera* | This work |
| mPAO1 pilD$^{12}$/pilA '+pBADGR'-*pilA*$^{chi}$ | 12 base duplication of residues 548-559 of *pilD, pilA ISphoA/hah insertion at position 163 in pilA, complemented with pilA-hxcT chimera* | This work |
| PAO1 PilS N323A | Hyperactive *pilS A867G* mutant | (2) |
| PAO1 PilS N323A | Hyperactive *pilS* mutant *A867G* complemented with pBADGR | This work |
| PAO1 *pilD*$^{12}$/ PilS N323A | Hyperactive *pilS A867G* mutant and 12 base duplication of residues 548-559 of *pilD* | This work |
| PAO1 *pilD*$^{12}$/ PilS N323A +pBADGr | Hyperactive *pilS A867G* mutant and 12 base duplication of residues 548-559 of *pilD* complemented with pBADGR | This work |
| PAO1 *pilT::Tn5* | ISphoA/hah insertion at position of 885 in *pilT* | (1) |
| PAO1 *pilD*$^{12}$/ *pilT::Tn5* | ISphoA/hah insertion in *pilT* 12 base duplication of residues 548-559 of *pilD* | This work |
| *E. coli strains* | | |
| DH5α | *F-φ80lacZΔM15 Δ(lacZYA-argF)U169 recA1 endA1 hsdR17(rk−, mk+) phoA supE44 thi-1 gyrA96 relA1 λ−* | Invitrogen |
| SM10 | *thi-1 thr leu tonA lacY supE recA::RP4-2-Tc::Mu (KmR)* | Invitrogen |
| Plasmids | | |
| pEX18Gm | Suicide vector for gene replacement | (3) |
| pBADGr | arabinose-inducible, broad host range complementation vector | (4) |
| pEX18Gm- *pilD*$^{12}$ | Construct to insert *pilD*$^{12}$ | This work |
| pEX18Gm- *pilD*$^{3}$ | Construct to insert *pilD*$^{3}$ | This work/IDT Gblock |
| pEX18Gm- *pilD6* | Construct to insert *pilD*$^{6}$ | This work/IDT Gblock |

| | | |
|---|---|---|
| pEX18Gm- *pilD*$^{9}$ | Construct to insert *pilD*$^{9}$ | This work/IDT Gblock |
| pEX18Gm- *pilD*$^{12var}$ | Construct to insert *pilD12*var | This work/IDT Gblock |
| pEX18Gm-PilB D388A | Construct to insert PilB D388A | This work |
| pBADGR-*pilA*$^{chi}$ | Construct to complement *pilA-hxcT chimera* | This work |
| pBADGr-*pilA* | Construct to complement *pilA* | This work |
| pBADGr-*pilB* | Construct to complement *pilB* | This work |
| pBADGr-PilB D388A | Construct to complement PilB D388A | This work |

**Supplementary Table S2. Primers used in this study**

| **Gene** | **Forward (5' → 3')** | **Reverse (5' → 3')** |
|---|---|---|
| Chromosomal knock in primers | | |
| *pilD*$^{12}$ | ACAAGAGCTCGAGTGGTGGCTTGCCG | CACAAAGCTTCAGCACGTCGTCGGC |
| *pilB* D388A | ACAAGAATTCATGAACGACAGCATCCAACTGAGC | ACAACCCGGGTTAATCCTTGGTCACGCGGTTGAC |
| Complementation primers | | |
| *pilA*$^{chi}$ | AATAGAATTCATGGATGTCGTGCAGTTCAGCTCCAGCCCGAAAGGACATCGGGGACAGCGGGGCTTTACCTTGATCGAA | TTGTTGGGCCCCGCATCGGCCTCTACGATG |
| *pilA* | AACAAGAGCTCCAAAGAGCTTGTTGCCGCG | TTGTTGGGCCCCGCATCGGCCTCTACGATG |
| *pilB* | ACAAGAATTCATGAACGACAGCATCCAACTGAGC | ACAACCCGGGTTAATCCTTGGTCACGCGGTTGAC |
| *pilB* D388A | ACAAGAATTCATGAACGACAGCATCCAACTGAGC | ACAACCCGGGTTAATCCTTGGTCACGCGGTTGAC |
